# AlphaGenome informed machine learning for pathogenicity prediction of germline mutations

**DOI:** 10.64898/2026.10.20.760593

**Authors:** Pankaj Kumar, Srishti Singh, Sandip Garai, Madhu Ramesh, Rangasai Chandra Goli, KC Nikhil, Tanmaya K. Sahu, Soumen Naskar, Vijai Pal Bhadana, Madhusudan R. Nandineni, KK Kanaka

**Affiliations:** ICAR- Indian Institute of Agricultural Biotechnology, Ranchi, India; Indian Institute of Technology, New Delhi, India; ICAR- Indian Agricultural Research Institute, New Delhi, India; Department of Biology, University of waterloo, Waterloo, ON, Canada; BRIC-Centre for DNA Fingerprinting and Diagnostics, Hyderabad, India

**Keywords:** Variant pathogenicity prediction, Machine learning, Functional genomics, AlphaGenome, Variant coverage, Clinical genomics

## Abstract

Accurate interpretation of germline variants remains a central challenge in clinical genomics. Existing *in silico* pathogenicity predictors often provide incomplete coverage or restricted to missense variants from the coding region, leaving the variants from splice sites, -UTR, and intronic regions under-scored. We present AGVarPred, a gradient-boosted classifier designed to predict the pathogenicity of germline mutants, with a broad consequence-class coverage across SNVs and indels. Here, the population allele frequency from gnomAD, functional consequence annotations from Ensembl VEP, and cell-type-specific functional genomic scores from the AlphaGenome platform were integrated to develop the classifier. Using a gene-level split of 134,002 high-confidence ClinVar variants across 9,222 genes, AGVarPred achieved an AUC-ROC of 0.9753 (95% CI 0.9731-0.9775) and a AUC-PR of 0.9606 on 12,385 scored internal test variants. A paired DeLong analysis showed that the full model significantly outperformed an allele-frequency-plus-VEP model (ΔAUC = 0.1539, 95% CI 0.1489-0.1589; p < 1 × 10^-300^), supporting a substantial incremental contribution from AlphaGenome-derived features. Performance remained strong when restricted to SNVs (AUC-ROC 0.9558) and was also high for indels (AUC-ROC 0.9532). On five external benchmarks, AUC-ROCs ranged from 0.528 on a functional-assay-derived benchmark to 0.977 on a clinically curated deafness database, reflecting its dependence on benchmark composition and label source. A separate analysis of the VIP Database, whose pathogenic labels are ClinVar-derived, showed consistent performance (AUC-ROC of 0.892) in held-out genes. AGVarPred scored 100% of evaluated variants in the external benchmarks, whereas AlphaMissense and REVEL covered only small fractions of non-missense variants. Model interpretation indicated that gnomAD allele frequency was the dominant individual predictor; the 111 AlphaGenome features provided complementary functional signal, with splicing and chromatin assays contributing to the largest share of the AlphaGenome SHAP mass. A web-based interface for AGVarPred is available at https://pankaj357.github.io/AGVarPred/ to facilitate the submission of normalized GRCh38 germline variants and retrieval of pathogenicity predictions using the frozen AGVarPred model.

## Introduction

The clinical interpretation of germline variants relies on computational predictors that aggregate sequence conservation, functional annotation, and population data into a single pathogenicity score. Tools such as CADD [1], REVEL [2], and AlphaMissense [3] have become widely used, but most were developed with a strong emphasis on missense variation and protein-sequence conservation. Other predictors, including protein sequence- and structure-informed approaches, are particularly effective for evaluating missense variants by modelling effects on protein function. Consequently, splice-region, synonymous, UTR, and intronic regulatory variants are often overlooked by these methods or receive less reliable scores. Because a major fraction of clinically relevant variation falls outside the canonical coding sequence, there is a need for predictors that can score variants across consequence-classes with a consistent model.

DNA foundation models are becoming vital in the functional genomics studies by learning sequence-to-function relationships directly from genomic context at scale. Unlike predictors focused primarily on protein-coding changes, platforms such as AlphaGenome can generate genome-wide, cell-type and assay-specific estimates of variant effects across regulatory, splicing, and coding regions. AlphaGenome enables a single framework to capture diverse molecular consequences, including changes in chromatin accessibility, transcription factor binding, gene expression, and RNA processing, that may underlie pathogenicity, particularly outside canonical missense variation. By providing functional predictions across hundreds of biosamples and assay types, AlphaGenome also offers a systematic way to prioritize context-dependent effects that would be difficult to infer from conservation or consequence annotation alone. Integrating these foundation-model-derived scores with established information such as population allele frequency and variant consequence could potentially improve pathogenicity prediction, particularly for all the variants other than missense variants (coding region).

At the same time, broader coverage introduces methodological challenges. Functional genomic scores are often correlated across tissues and assays, complicating model interpretation. Population allele frequency is itself an extremely strong predictor of benign status, and models that rely heavily on it, may generalize poorly to settings where allele frequency is uninformative or unavailable. Benchmarking must also account for the fact that different tools cover different variant subsets and the comparisons restricted to common subsets can be biased towards the strengths of the most limited tool. Finally, many public benchmarks overlap with the training data of existing predictors, making the independent evaluation difficult. Here we describe AGVarPred, a LightGBM-based [5] pathogenicity predictor trained on a gene-held-out ClinVar [6] gold standard and evaluated on multiple independent benchmarks. We emphasize conservative interpretation and report its performance across diverse benchmarks, dissect the contribution of allele frequency versus functional scores, evaluate coverage explicitly, and discuss limitations including dependence on AlphaGenome. Further we have also discussed the restriction of the current evaluation to variants represented in the final feature matrices, and limited transportability of calibration across external benchmarks. Our goal is to establish whether a single model can provide strong discrimination across diverse variant classes and provide probability estimates that are well calibrated within the development setting, while explicitly evaluating their transportability across external benchmarks rather than to claim universal superiority over specialized missense predictors.

## Methods

### Study Overview

AGVarPred is a germline variant pathogenicity predictor evaluated across single-nucleotide variants (SNVs) and indels mostly focusing on broad consequence-class coverage. It integrates cell-type-specific functional genomic scores from the AlphaGenome platform with population allele frequencies from gnomAD and functional consequence annotations from Ensembl Variant Effect Predictor (VEP). The work was motivated by the observation that many existing *in silico* predictors are optimized for missense variation and provide incomplete coverage of splice, synonymous variants of coding region, Untranslated region (UTR), and intronic variants. The AlphaGenome-derived feature set used in this study was applied to the variants supported by the feature-extraction pipeline, allowing evaluation across multiple consequence classes.

The development workflow proceeded in four stages. First, a high-confidence ClinVar-derived gold standard was assembled, filtered to germline variants with explicit expert review, and partitioned at the gene level into training, calibration, and test sets. Second, each variant was annotated with AlphaGenome functional scores, gnomAD allele frequency, and VEP-derived features, and the resulting high-dimensional feature matrix was reduced by nested feature elimination. Third, a regularized gradient-boosted tree classifier was optimized with gene-grouped cross-validation, calibrated with isotonic regression on the held-out calibration set, and threshold-tuned to maximize F1. Fourth, the frozen pipeline was evaluated on the held-out ClinVar test set and on five external benchmarks. Its prediction performance was compared with CADD, AlphaMissense, and REVEL using coverage-aware evaluation schemes, DeLong testing, and decision-curve analysis. Model interpretation was performed with SHAP values aggregated based on tissue and assay type by dissecting the performance across variant consequence classes. All code, data manifests, and model artifacts required to inspect and reproduce the reported analysis are provided in the accompanying repository and Zenodo archive; reproduction of AlphaGenome-dependent feature extraction requires access to the AlphaGenome platform. AGVarPred was evaluated across SNVs and indels in the present study and requires AlphaGenome functional scores as input; users without a gnomAD VCF can use the no-allele-frequency model; AlphaGenome-derived features remain required for this model. An overview of the AGVarPred workflow is shown in Figure 1.

**Figure 1.**
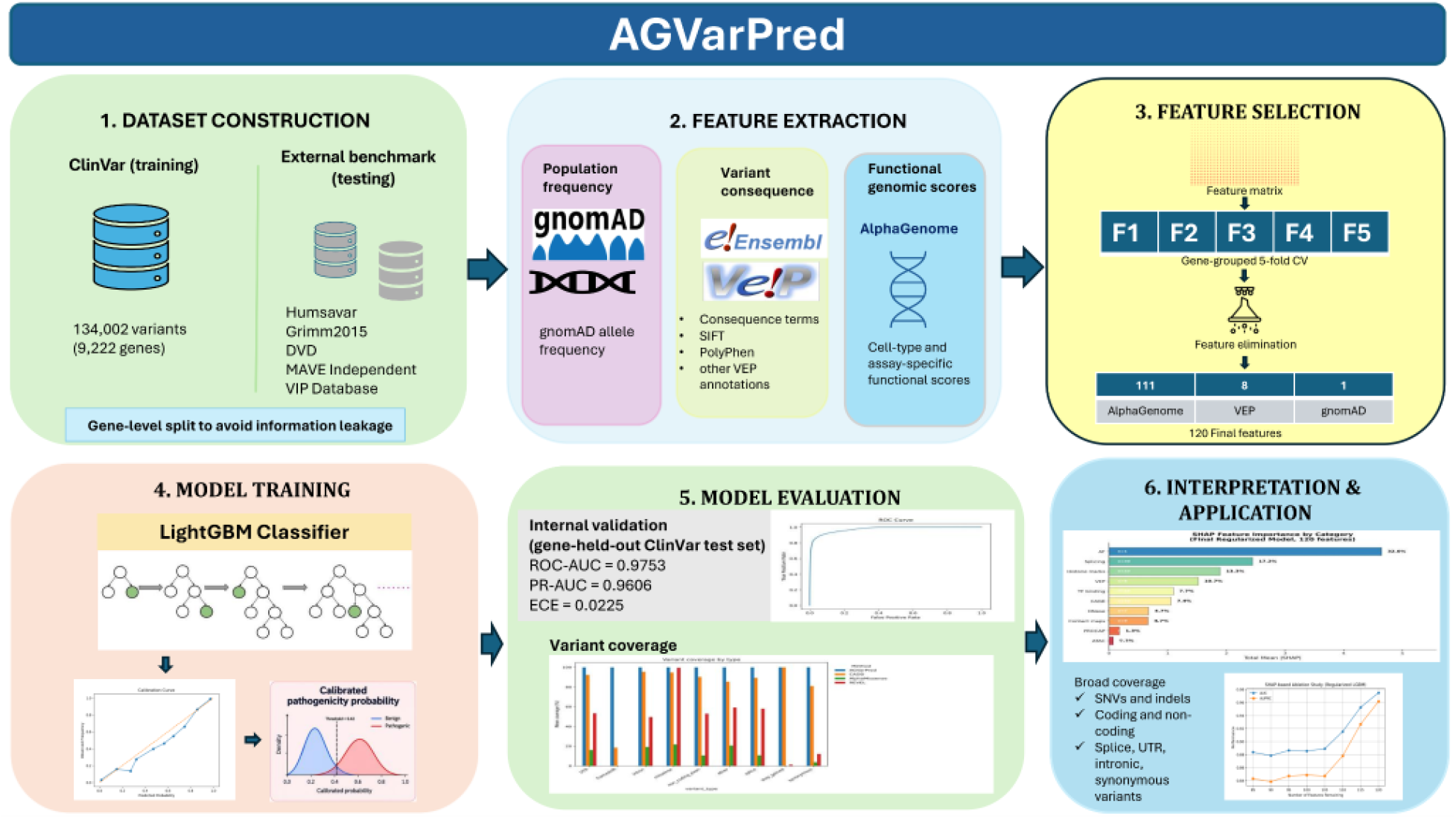
AGVarPred workflow overview. Overview of the AGVarPred pipeline. (1) ClinVar variant_summary.txt was filtered to a 134,002-variant high-confidence gold standard. (2) Variants were split at the gene level into 80% training, 10% calibration, and 10% test sets. (3) Features were extracted with the AlphaGenome SDK, gnomAD allele frequency, and Ensembl VEP. (4) Nested feature elimination reduced the panel to 120 features. (5) A LightGBM classifier was trained, isotonically calibrated, and threshold-tuned (0.42). (6) The frozen pipeline was evaluated on the scored internal ClinVar test subset and on five external benchmarks; the VIP Database was analyzed separately as a ClinVar-held-out-gene consistency analysis.

### Dataset construction

The primary training labels were obtained from the NCBI ClinVar variant_summary.txt flat file (release 31 March 2024), which aggregates submitted interpretations for human germline variants. Variants were retained only if they were annotated on the GRCh38 genome assembly, carried an OriginSimple value of germline, and satisfied one of the three highest review-status tiers: criteria provided, multiple submitters, no conflicts; reviewed by expert panel; or practice guideline. Only records with an unambiguous clinical significance of Pathogenic or Benign were kept, and any record containing the string conflicting was excluded to avoid ambiguous labels. Rows missing any of the four VCF coordinates (chromosome, position, reference allele, alternate allele) were dropped, and the remaining variants were deduplicated by chrom:pos:ref:alt.

The resulting ClinVar gold-standard dataset comprised 134,002 unique variants in 9,222 genes, of which 40,584 (30.3%) were pathogenic and 93,418 (69.7%) benign (Table 1). The dataset contained multiple variant types; SNVs were the majority, but deletions, duplications, insertions, indels, microsatellites, and other variant classes were also present. Feature extraction, model training, calibration, and internal evaluation were performed only on the variants successfully represented in the final feature matrices: 86,378 training-set variants, 13,201 calibration-set variants, and 12,385 test-set variants. The build script did not apply any phenotype, disease, or allele-frequency filters beyond the germline, review-status, and label filters described above, so the initial dataset represents a broad, unselected sample of high-confidence ClinVar interpretations.

**Table 1.** ClinVar gold-standard dataset and gene-level splits.

| Dataset | Variants | Genes | Pathogenic | Benign |
| --- | --- | --- | --- | --- |
| ClinVar gold standard | 134,002 | 9,222 | 40,584 (30.3%) | 93,418 (69.7%) |
| Training (initial split) | 101,604 | 7,377 | 27,788 (27.3%) | 73,816 (72.7%) |
| Calibration (initial split) | 16,223 | 922 | 6,537 (40.3%) | 9,686 (59.7%) |
| Test (initial split) | 16,175 | 923 | 6,259 (38.7%) | 9,916 (61.3%) |
| Training (scored feature matrix) | 86,378 | — | 26,896 (31.1%) | 59,482 (68.9%) |
| Calibration (scored feature matrix) | 13,201 | — | 5,522 (41.8%) | 7,679 (58.2%) |
| Test (scored feature matrix) | 12,385 | — | 4,576 (36.9%) | 7,809 (63.1%) |

Note. The initial gene-level splits are the full train/calibration/test partitions; the scored rows show the variants represented in the final feature matrices on which training, calibration, and internal evaluation were performed. Gene counts are not reported for the scored subsets because the matrices are keyed by variant, not by gene.

To prevent information leakage, the dataset was split at the gene level rather than at the variant level. All unique gene symbols were first randomly divided into an 80% training partition and a 20% temporary partition using sklearn.model_selection.train_test_split with random_state=42. The temporary partition was then split again into 10% calibration and 10% test sets with the same random seed. Because every gene appeared in exactly one partition, the resulting sets contained no shared genes. The training set contained 101,604 variants in 7,377 genes (27,788 pathogenic, 73,816 benign), the calibration set 16,223 variants in 922 genes (6,537 pathogenic, 9,686 benign), and the test set 16,175 variants in 923 genes (6,259 pathogenic, 9,916 benign). The class distributions were similar across splits (training 27.3% pathogenic, calibration 40.3% pathogenic, test 38.7% pathogenic), with the calibration and test sets being modestly enriched for pathogenic variants because of the random gene-level split. Split sizes and class counts, together with the scored subsets represented in the final feature matrices, are summarized in Table 1.

### Leakage prevention

Gene-level partitioning was the central leakage-control strategy. By ensuring that no gene was present in more than one of the training, calibration, or test sets, we prevented the model from learning gene-specific effects, such as known loss-of-function intolerance, regional hypermutability, or variant-hotspot positions that would inflate performance on held-out variants from the same genes. This design also mirrors realistic use, in which a variant may be encountered in a gene that was seen during training only through different variants. A purely variant-level split would have allowed different variants in the same gene to appear in both training and testing, producing optimistic estimates for genes with strong pathogenic or benign clustering.

The gene-grouping principle was enforced during both feature selection and hyperparameter optimization. Nested feature elimination used GroupKFold (n_splits=5) from scikit-learn [7], with the grouping variable set to gene, so that every training fold trained on one set of genes and validated on a disjoint set. Likewise, the Optuna hyperparameter search used five gene-group folds, and the objective function was the mean area under the precision-recall curve (AUPRC) across these folds. Calibration and threshold optimization were performed exclusively on the held-out calibration genes, which were also absent from the training set.

External benchmarks were required to satisfy independence criteria before inclusion: zero variant overlap with the ClinVar train/cal/test sets, zero gene overlap with the ClinVar train/cal/test sets, no ClinVar-derived labels, and germline-only variation. These criteria were verified with dedicated audit scripts that compared chrom:pos:ref:alt coordinates and HGNC gene symbols against the training set. Several candidate benchmarks were excluded because they violated one or more of these criteria. A ClinVar 3-star holdout benchmark was removed after an audit revealed 91.6% gene overlap and 39% variant overlap with the training data. Other excluded datasets included somatic cancer-hotspot variants, phenotype-derived de novo case/control sets, constrained-gene panels with hundreds of genes already present in training, and MAVE assays for genes such as BRCA1 and PTEN that were present in the training set. The retained benchmarks and their independence audits are summarized in Supplementary Note 2. A JSON file (train_gene_set.json) recording the 9,222 genes used for training, calibration, and testing is distributed with the repository so that future benchmarks can be checked for independence. We note that these independence criteria do not guarantee independence from the training data of the comparison tools. Public benchmarks such as Humsavar have been used extensively in the development of CADD, REVEL, and other predictors, and some label circularity with those tools is likely unavoidable.

### Variant annotation

Functional genomic features were generated with the AlphaGenome Python SDK (version 0.6.1). For each gene, input variant lists were prepared in VCF-like tabular format (chromosome, position, reference allele, alternate allele, gene symbol) and processed with the SDK’s variant scorers using the recommended variant-scorer collection and a 1 Mb reference interval centered on each variant. Each scored variant produced a vector of raw scores, with one column per assay track named {output_type} {biosample_name} {track_name}. The output matrices were written as Apache Parquet files and then merged across all genes and all scoring runs. The SDK calls require valid API keys, which are supplied via the ALPHAGENOME_API_KEYS environment variable as a comma-separated list. AlphaGenome scores are currently available only through this SDK; reproducibility therefore depends on continued access to the platform.

Population allele frequency and VEP annotations were extracted from the gnomAD [8] exomes release 2.1.1 liftover to GRCh38 (gnomad.exomes.r2.1.1.sites.liftover_grch38.vcf.bgz) using pysam.VariantFile. The allele-frequency field AF was read for each matched variant; when absent, the allele frequency was set to zero. VEP annotations were taken from the CSQ INFO field, which contains a comma-separated list of consequence annotations across Ensembl transcripts. The VEP string was parsed for the first transcript listed, and the following fields were extracted: SIFT prediction and score, PolyPhen prediction and score, consequence term, IMPACT level, Loss-of-Function (LoF) flag, protein position, and amino-acid change. These annotations were generated by gnomAD using Ensembl VEP [9] release 85 (GENCODE v19); no separate VEP re-annotation was performed.

Variants were identified consistently throughout the pipeline with a variant_id of the form chr{chrom}_{pos}_{ref}_{alt}. Chromosome identifiers were normalized to include the chr prefix during feature labeling. During feature labeling, the ClinVar labels and gene symbols were joined to the AlphaGenome matrices by this identifier, and the gnomAD AF and VEP-derived fields were appended. The merged matrices were then written to per-split Parquet datasets (final_dataset_parts_train, final_dataset_parts_cal, final_dataset_parts_test) that served as input to feature selection and model training. For external benchmarks, VEP annotations were preprocessed to match the training pipeline’s encoding, including one-hot encoding of vep_Consequence and ordinal encoding of SIFT, PolyPhen, IMPACT, and LoF categories. A previous preprocessing omission of one-hot consequence encoding was corrected before the final analyses; all reported external results reflect the corrected encoding.

### Feature engineering

The merged feature matrix contained 4706 columns: one per AlphaGenome assay track, plus derived VEP and allele-frequency fields. Before modeling, features were separated into numeric and categorical types. Numeric features were imputed with the median computed within each training fold during feature selection, and with the global training-set median for the final pipeline. Median imputation was chosen because it is robust to skewed distributions and because the AlphaGenome score matrix contains many missing values for tracks in which a given variant class cannot be scored. All numeric features were scaled with sklearn.preprocessing.RobustScaler, which centers on the median and scales by the interquartile range and is therefore insensitive to allele-frequency outliers and extreme functional scores.

VEP-derived variables were represented as numeric model inputs after preprocessing. No categorical columns were passed directly to LightGBM. The final selected VEP features comprised vep_IMPACT_score, vep_SIFT_score, vep_PolyPhen_score, vep_LoF, vep_is_synonymous, and one-hot consequence indicators for synonymous, intronic, and downstream-gene variants. Missing numeric values were handled by median imputation using training-derived medians.

Additional derived features included log10_gnomAD_AF (computed as log10 of gnomAD_AF + 1e-8), a binary AF_missing flag, an is_ultra_rare flag for alleles with frequency below 0.01%, and binary indicators for missense, synonymous, stop-gained, frameshift, splice-region, and high-confidence loss-of-function consequences. These flags allowed the model to learn both the magnitude of allele frequency and the presence or absence of canonical consequence categories.

Feature selection used nested feature elimination driven by LightGBM gain importance. On each of five gene-group folds, the full feature matrix was preprocessed without leakage, a LightGBM classifier with fixed base hyperparameters (300 trees, seed 42, gain importance) was fit, and the top n features by gain importance were selected. A reduced model was then retrained on those n features and evaluated on the held-out fold. This procedure was repeated for n in {40, 60, 80, 100, 120, 140, 160, 200, 250, 300, 400, 500, 750, 1000, 1500, 2000}. The optimal panel size was chosen using a one-standard-deviation rule applied to mean cross-validated AUPRC: the smallest n whose mean AUPRC was within one standard deviation of the best observed mean AUPRC. This rule was preferred over the raw maximum because it favors parsimony and reduces the risk of overfitting to noisy importance ranks, while retaining nearly all of the cross-validated predictive performance (Supplementary Figure S1).

The final selected panel contained 111 AlphaGenome functional-score features, one gnomAD allele-frequency feature (gnomAD_AF), and eight VEP-derived features (vep_IMPACT_score, vep_SIFT_score, vep_PolyPhen_score, vep_LoF, vep_is_synonymous, vep_Consequence_synonymous_variant, vep_Consequence_intron_variant, vep_Consequence_downstream_gene_variant) (Supplementary Table S1). A log10_gnomAD_AF feature was derived during preprocessing but was not retained in the final selected panel. The selected feature panel was supplied to LightGBM as numerical model inputs. A secondary no-allele-frequency model was trained analogously after removing the single AF feature, yielding a 119-feature panel and a calibrated threshold of 0.45 (Supplementary Figure S6).

### Model development

The classifier used was a gradient-boosted decision-tree ensemble implemented in LightGBM (LGBMClassifier). Hyperparameters were optimized with Optuna [10] (Tree-structured Parzen Estimator sampler, seed 42) over 100 trials. The objective was the mean five-fold gene-group AUPRC on the training set. The search space was: n_estimators 200-500; learning_rate 0.01-0.08; num_leaves 20-80; max_depth 3-8; min_child_samples 10-100; subsample 0.6-1.0; colsample_bytree 0.6-1.0; reg_alpha 1e-3-10.0 (log scale); and reg_lambda 1e-3-10.0 (log scale) (Supplementary Table S2). The relatively narrow ranges and emphasis on regularization parameters were chosen to avoid overfitting to the large but correlated AlphaGenome feature panel.

Class imbalance was addressed using scale_pos_weight = 2.2116, calculated once from the full training set (59,482 benign / 26,896 pathogenic) and held fixed across all cross-validation folds and the final model fit. No early stopping was used; the search explicitly preferred shallow trees and regularization to reduce overfitting. No class rebalancing by resampling was applied. The best trial was trial 64, which achieved a cross-validated AUPRC of 0.9676. The chosen hyperparameters were: n_estimators 485; learning_rate 0.0653; num_leaves 73; max_depth 8; min_child_samples 91; subsample 0.6013; colsample_bytree 0.6757; reg_alpha 0.0373; reg_lambda 0.0140; scale_pos_weight equal to the benign-to-pathogenic ratio; random_state 42; and n_jobs -1.

The final model was trained on the full training set using the 120 selected features. Training-set medians and a RobustScaler were fit on the training data and applied to the calibration and test sets. The model was frozen after training; no weight updates were performed during calibration or external validation. The same frozen pipeline was used for all internal and external scoring, including the no-allele-frequency pipeline for benchmarks where gnomAD AF was unavailable.

### Probability calibration

Raw predicted probabilities from the LightGBM classifier were calibrated with sklearn.isotonic.IsotonicRegression(out_of_bounds=’clip’). Isotonic regression was chosen because it is non-parametric and can correct the typically under-confident or over-confident outputs of gradient-boosted classifiers without assuming a specific parametric form. The calibrator was fit exclusively on the scored calibration set, comprising 13,201 variants represented in the final feature matrix and originating from 922 genes not used for training. The calibrated probabilities were then used for internal evaluation and threshold selection.

The binary operating threshold was selected on the calibrated probabilities from the scored calibration set (13,201 variants; 41.8% pathogenic). Thresholds from 0.05 to 0.95 were evaluated in steps of 0.01, and the threshold maximizing F1 score on the calibration set was retained. Thus, 0.42 represents an empirically selected operating threshold for this ClinVar-derived development distribution rather than a clinically established or gold-standard pathogenicity cutoff. For the full model this threshold was 0.42; for the no-allele-frequency model it was 0.45. Because this threshold was optimized on a ClinVar-derived calibration set with substantially higher pathogenic prevalence than several external benchmarks, it was not assumed to be optimal for other populations or clinical settings.

The final inference pipeline therefore comprised: median imputation using training-set medians; RobustScaler standardization; LightGBM probability prediction; isotonic calibration; and classification using the calibrated 0.42 threshold. The pipeline was serialized as final_pipeline.pkl and used without modification for all internal and external evaluations. Calibration quality was assessed on the internal ClinVar test set and externally using Brier score, 10-bin ECE, calibration intercept, and calibration slope on each benchmark.

### Internal validation

Internal performance was evaluated on the 12,385 test-set variants represented in the final feature matrices. The following metrics were computed from the calibrated probabilities and the fixed 0.42 threshold: area under the receiver-operating-characteristic curve (ROC-AUC), area under the precision-recall curve (PR-AUC), Matthews correlation coefficient (MCC), precision, recall, F1 score, Brier score, and expected calibration error (ECE). ECE was computed with ten equal-width probability bins as the sum of absolute differences between bin-wise observed pathogenic frequency and mean predicted probability, weighted by bin size. The confusion matrix was also saved.

On the test set, AGVarPred achieved ROC-AUC 0.9753 (95% bootstrap confidence interval 0.9731-0.9775), PR-AUC 0.9606, F1 0.8945, precision 0.8952, recall 0.8938, MCC 0.8327, Brier score 0.0568, and ECE 0.0225. The bootstrap confidence interval for ROC-AUC was derived from 2,000 stratified resamples of the test set with replacement. A stratified dummy classifier achieved an AUC of 0.4959, confirming that the observed discrimination was not driven by class imbalance.

To assess robustness across molecular contexts, test-set variants were classified by VEP consequence into missense, synonymous, stop-gained, frameshift, splice, UTR, intron, non-coding exon, and other. Per-class AUCs for evaluable classes were: missense 0.9842, stop-gained 1.000, frameshift 0.9983, splice 0.9997, UTR 1.000, intron 0.9995, and non-coding exon 1.000. The synonymous class contained no pathogenic variants in the test set, so AUC could not be computed; a large ‘other’ category (7,352 variants, 3,668 pathogenic) had an AUC of 0.9413. These results indicate that the model discriminated pathogenic from benign variants consistently across evaluable consequence classes on the internal ClinVar test set. External per-class performance was more variable and is reported in the Results.

### External validation

The frozen AGVarPred pipeline was evaluated on five independent benchmarks selected to cover diverse variant types, curation sources, and disease contexts (Table 2). Every benchmark satisfied independence from the ClinVar training data: no overlapping variants and no overlapping genes. Benchmark-specific build scripts performed coordinate liftover where necessary, filtered for non-training genes, and removed variants present in the ClinVar splits.

**Table 2.** External benchmark summary.

| Benchmark | Total variants | Pathogenic | Benign | Genes | Scored by AGVarPred |
| --- | --- | --- | --- | --- | --- |
| UniProt Humsavar | 14,718 | 1,317 | 13,401 | 5,681 | 14,718 |
| MAVE Independent | 7,557 | 3,751 | 3,806 | 25 | 7,557 |
| gnomAD Common Benign | 4,989 | 0 | 4,989 | 585 | 4,989 |
| Grimm et al. 2015 Selected | 5,255 | 512 | 4,741 | 3,112 | 5,253 |
| DVD (Deafness Variation Database) | 8,388 | 195 | 8,193 | 12 | 8,388 |
| VIP Database (consistency analysis only) | 3,880 | 1,075 | 2,805 | 904 | 3,880 |

Note. The five external benchmarks (top five rows) satisfy gene-holdout and variant-holdout independence from the ClinVar training data and have labels derived independently of ClinVar. The VIP Database (bottom row) is reported separately as a ClinVar-held-out-gene consistency analysis because all 1,075 of its pathogenic labels are derived from overlapping ClinVar annotations. Two Grimm2015 benchmark variants were not scored because they mapped to the unsupported alternate contig chr19\_KI270938v1\_alt; all reported performance metrics therefore use the 5,253 scored variants.

**UniProt Humsavar** [11] is a manually curated set of human missense variants with disease-association labels. After restricting to non-training genes and removing variants present in the ClinVar splits, the benchmark contained 14,718 variants (1,317 pathogenic, 13,401 benign). The Humsavar benchmark initially contained 15,185 variants; 467 variants (454 on chrX, 9 on chrY, and 4 with non-ACGTN alleles) were excluded because a chromosome-name casing error in the benchmark preprocessing pipeline caused AlphaGenome to reject sex-chromosome contigs. The evaluated set therefore comprised 14,718 variants. Of the excluded variants, 208 were pathogenic. AGVarPred scored all 14,718 variants and achieved ROC-AUC 0.826 (95% CI 0.817-0.835), PR-AUC 0.275, and MCC 0.295 (95% CI 0.277-0.311). On the common subset scored by all tools (4,631 variants), AGVarPred AUC was 0.833, compared with 0.900 for CADD, 0.883 for AlphaMissense, and 0.943 for REVEL.

**MAVE Independent** was built from MaveDB [12] multiplexed assay scores for 25 assay targets (18 genes and 7 enhancer/construct targets) absent from the training set. Within each target, variants in the bottom quintile of the assay-score distribution were labeled “low-function” and variants in the top quintile were labeled “high-function”. These labels represent molecular assay function rather than clinical pathogenicity. AGVarPred scored 7,557 variants and achieved ROC-AUC 0.528 (95% CI 0.514-0.541), PR-AUC 0.511, and MCC 0.051 (95% CI 0.026-0.073). All compared tools performed near-random on this benchmark (CADD 0.562, AlphaMissense 0.697, REVEL 0.699 on their respective scored sets). These results indicate that molecular functional assay readouts in this benchmark are not equivalent to clinical pathogenicity labels; a variant can severely disrupt molecular function without being classified as pathogenic in the clinic, and vice versa. The MAVE benchmark also contained several assay targets that were not conventional gene-level identifiers; these were retained as provided by the source dataset and should be interpreted as assay targets rather than canonical gene entities.

**gnomAD Common Benign** consisted of common exonic single-nucleotide variants (allele frequency >0.1%) from non-training genes, all labeled benign. Because there were no pathogenic variants, ROC-AUC and MCC were not applicable; the benchmark was evaluated with false-positive rate (FPR) and accuracy. AGVarPred scored 4,989 variants with an FPR of 0.0024 (95% CI 0.001-0.004) and accuracy 0.9976.

**VIP Database (consistency analysis only)** contained clinical-lab germline variants from Baylor College of Medicine [13]. Benign labels corresponded to the VIP LB category, while pathogenic labels corresponded to native Path, Lpath, or Path_NR categories, or review entries cross-referenced to ClinVar Pathogenic/Likely pathogenic. Because all 1,075 pathogenic labels were derived from overlapping ClinVar annotations, VIP was not treated as an independent external benchmark; instead, it was used to assess consistency of ClinVar-derived classifications in held-out genes. After GRCh37-to-GRCh38 liftover and removal of training-set genes and variants, 3,880 variants remained (1,075 pathogenic, 2,805 benign across 904 genes). AGVarPred achieved ROC-AUC 0.892 (95% CI 0.881-0.903), PR-AUC 0.753, and MCC 0.507 (95% CI 0.484-0.529).

**Grimm et al. 2015** selected [14] combined the VariBenchSelected, predictSNPSelected, and SwissVarSelected datasets originally assembled to avoid circularity with predictor training sets. After additional filtering for non-training genes, variant deduplication, and GRCh37-to-GRCh38 liftover, 5,253 scored variants (512 pathogenic, 4,741 benign) from 5,255 total variants. AGVarPred achieved ROC-AUC 0.745 (95% CI 0.727-0.762), PR-AUC 0.188, and MCC 0.233 (95% CI 0.210-0.257). On the jointly scored subset with REVEL (5,183 variants), AGVarPred outperformed REVEL (AUC 0.746 versus 0.641, DeLong q < 0.001).

**DVD (Deafness Variation Database)** [15] contained expert-curated variants associated with deafness and hearing loss. After gene and variant holdout, 8,388 variants remained (195 pathogenic, 8,193 benign across 12 genes). AGVarPred achieved ROC-AUC 0.977 (95% CI 0.967-0.985), PR-AUC 0.788, and MCC 0.258 (95% CI 0.239-0.275).

Across the five independent benchmarks, AGVarPred achieved full-coverage ROC-AUCs of 0.826 on Humsavar, 0.528 on MAVE Independent, 0.745 on Grimm2015, 0.977 on DVD, and an FPR of 0.0024 on gnomAD Common Benign. The separate VIP consistency analysis achieved ROC-AUC 0.892. Three comparison modes were used when benchmarking against CADD, AlphaMissense, and REVEL. Full-coverage evaluation reported each tool on every variant it could score, reflecting real-world usability but allowing coverage to differ. Pairwise common-subset evaluation compared AGVarPred with each competitor only on variants scored by both tools, balancing fairness with sample size. All-tools common-subset evaluation required all four tools to have a score and was reported where informative. Because AlphaMissense and REVEL are missense-biased, their coverage dropped sharply for splice, synonymous, and non-coding variants; CADD phred scores were available for most variants but are not probabilistically calibrated. Standard published thresholds were used for competitors: CADD phred ≥15, AlphaMissense ≥0.5, and REVEL ≥0.5. Confidence intervals for all external metrics were computed from 1,000 bootstrap resamples.

### Statistical analysis

Confidence intervals were calculated with the percentile bootstrap. For the internal test set, 2,000 stratified resamples with replacement were used to obtain the 95% confidence interval for ROC-AUC. For external benchmarks, 1,000 unstratified bootstrap resamples with replacement were used to obtain confidence intervals for ROC-AUC and MCC when both classes were present and the evaluated subset contained at least 10 variants. Very small subsets (n < 10) and single-class subsets were excluded from bootstrap confidence intervals.

Pairwise statistical comparisons of correlated ROC curves were performed with DeLong’s test [16]. For each benchmark, AGVarPred was compared separately with CADD, AlphaMissense, and REVEL on the set of variants scored by both tools. The test statistic was the standardized difference in AUC estimates, with standard errors derived from the joint covariance of the two ROC curves. Raw p-values were adjusted across all pairwise comparisons using the Benjamini-Hochberg false-discovery-rate procedure [17], and significance was reported at q < 0.05. The correction was applied to the set of valid pairwise DeLong tests across the five independent benchmarks and three competitors (11 valid pairwise DeLong tests in total: 3 on Humsavar, 3 on MAVE Independent, 2 on Grimm2015 because the AlphaMissense subset contained no pathogenic variants, and 3 on DVD; gnomAD Common Benign was excluded because it contains only benign variants, and VIP was excluded because its pathogenic labels are ClinVar-derived and analyzed separately as a consistency analysis). Because AlphaMissense and REVEL cover only small fractions of non-missense variants, some pairwise comparisons had insufficient sample size or single-class subsets and were marked not applicable. We caution that common-subset comparisons are inherently biased toward the variant types that the most restrictive competitor can score.

A paired DeLong test was additionally used to compare the full AGVarPred model with the allele-frequency-plus-VEP model on the same internal test variants. This analysis evaluated whether incorporation of AlphaGenome-derived features provided a statistically significant improvement in discrimination while accounting for the correlation between predictions generated on the same variants. The difference in ROC-AUC and its 95% confidence interval were calculated from the paired ROC curves.

Decision-curve analysis [18] was performed to evaluate the potential clinical utility of probabilistic predictions. On the internal test set, AGVarPred produced positive net benefit from threshold 0.01 through 0.99 and exceeded both treat-all and treat-none strategies across the same range. On external benchmarks, AGVarPred exceeded both reference strategies on Humsavar for thresholds 0.01-0.20, on Grimm2015 for 0.02-0.17, and on DVD for 0.01-0.03; it produced near-zero net benefit on gnomAD Common Benign by correctly classifying almost all benign variants. Net benefit was computed at thresholds from 0.01 to 0.99 in steps of 0.01 as 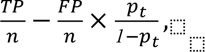, where TP and FP are true and false positives, n is the total number of variants, and p_t is the decision threshold. Net-benefit curves for AGVarPred were plotted against the “treat all” and “treat none” reference strategies. Because CADD reports phred scores rather than calibrated probabilities, CADD was excluded from the decision-curve comparison; AlphaMissense and REVEL were included on their jointly scored subsets with AGVarPred. Integrated net benefit, maximum net benefit, and the threshold ranges over which AGVarPred produced positive net benefit and exceeded both treat-all and treat-none strategies were summarized for each benchmark.

### Model interpretation

Model interpretation was based on SHAP (SHapley Additive exPlanations) values computed with shap.TreeExplainer [19] on a random sample of 5,000 training-set variants. SHAP values were computed from the uncalibrated LightGBM model output and are therefore expressed on the model’s log-odds margin scale rather than as differences in calibrated pathogenic probabilities. For each selected feature, the mean absolute SHAP value across the sample was used as an importance score. A global summary plot and a ranked importance table were generated and saved as supplementary material. Because the 111 AlphaGenome features encode cell-type and assay type in their names, they were parsed into three components: output type (assay), biosample name, and track identifier. Assay types included CAGE, DNase I hypersensitivity, ATAC-seq, TF ChIP-seq, histone ChIP-seq, splice-site usage, splice junctions, splice sites, chromatin contact maps, and PRO-cap. Biosamples were mapped heuristically to 13 broad tissue groups: brain/CNS, liver/digestive, heart/vasculature, blood/immune, reproductive, embryonic/stem, lung, kidney/urinary, eye/sensory, skin, muscle, connective/mesenchyme, and other. SHAP importance was then aggregated by tissue group and by assay type. Bootstrap confidence intervals were computed by resampling features within each group. The brain/CNS and liver/digestive groups contributed the largest cumulative SHAP importance among AlphaGenome features (approximately 22% and 20% of the AlphaGenome SHAP mass, respectively), followed by heart/vasculature (∼9.8%) and blood/immune (∼8.4%). Splicing and transcriptomic assays accounted for a large share of the aggregated AlphaGenome contribution.

We emphasize that these SHAP-based interpretations are descriptive, not causal. The dominant individual predictor was gnomAD allele frequency (mean absolute SHAP 4.648), followed by VEP IMPACT score (0.437); the top AlphaGenome feature contributed only 0.290. Many AlphaGenome tracks are correlated across biosamples and assays, and the mapping of biosamples to tissue groups is heuristic. Tissue and assay rankings should therefore be interpreted as reflecting which signals were available and selected by the model, not as evidence that pathogenicity is driven by specific tissues or assays in a biological sense.

Variant-type interpretation used VEP consequence flags and terms to assign each variant to one of nine classes. Assignment followed a fixed priority order such as stop-gained, frameshift, splice, missense, synonymous, UTR, intron, non-coding exon, other so that each variant received a single, mutually exclusive label. Performance metrics were computed within each class, and a non-missense comparison was constructed by evaluating AGVarPred and competitors only on variants that were not missense. Because competitor coverage differs by variant type, the non-missense comparison was performed on the AGVarPred-scored universe to avoid coverage-based artifacts. AGVarPred maintained non-missense AUCs above 0.68 on Humsavar and above 0.92 on DVD, whereas AlphaMissense and REVEL frequently could not score these variants.

### Software availability

AGVarPred is distributed as an installable Python package with a command-line interface and a Python API. The package can be installed from the GitHub repository with pip install -e . and is compatible with Python 3.11, 3.12, and 3.13. The CLI supports subcommands including predict, list-models, and --version. A typical prediction command accepts a VCF file, an output path, and either a directory of precomputed AlphaGenome features or a local gnomAD VCF; if the gnomAD VCF is unavailable, the package automatically falls back to the bundled no-allele-frequency model. The Python API exposes AGVarPredAutoPredictor, FeatureGenerator, and lower-level feature-engineering classes in the agvarpred_core module.

The source code is hosted at https://github.com/pankaj357/AGVarPred and released under the Apache 2.0 license. A frozen manuscript archive, including trained model artifacts, ClinVar splits, and external benchmark data, is available on Zenodo at https://doi.org/10.5281/zenodo.20955782. Model directories contain manifest YAML files with model version, training date, feature count, threshold, calibration method, and SHA256 checksums for reproducibility. The full model manifest reports 120 selected features, isotonic calibration, a robust scaler, median imputation, and a 0.42 threshold.

The repository includes a comprehensive test suite run with pytest, covering feature engineering, VEP parsing, gnomAD querying, model loading, CLI behavior, and predictor output. Continuous integration is configured through GitHub Actions. The full research reproducibility pipeline, including data preparation, feature extraction, feature selection, model training, ablation studies, and external validation, is provided in the AGVarPred-training/ directory. Environment variables ALPHAGENOME_API_KEYS (comma-separated API keys) and GNOMAD_VCF (path to the gnomAD VCF) are required to re-run feature extraction. A pinned dependency list and a reproducibility checklist are included in the repository to facilitate independent replication of the training and evaluation workflow. The package is formatted with black, versioned with semantic version tags, and includes a CITATION.cff file and a LICENSE file (Apache 2.0). Documentation covering installation, API usage, and reproducibility is provided in the docs/ directory. Example VCF and precomputed AlphaGenome feature files in examples/ allow users to run a minimal prediction without external API access. All scripts that previously contained hardcoded API keys were refactored to read keys from the ALPHAGENOME_API_KEYS environment variable, and a credential scan confirmed no remaining hardcoded secrets in the current working tree. Reproducibility of AlphaGenome-dependent feature extraction therefore depends on continued access to the AlphaGenome platform and the applicable provider terms.

A web-based interface for AGVarPred is available at https://pankaj357.github.io/AGVarPred/ to facilitate the submission of normalized GRCh38 germline variants and retrieval of pathogenicity predictions using the frozen AGVarPred model. The web application uses a React-based frontend and a FastAPI backend and integrates AlphaGenome-derived functional genomic features with gnomAD allele-frequency and Ensembl Variant Effect Predictor (VEP) annotations. The service provides calibrated pathogenicity scores and corresponding classifications using the model’s predefined operating threshold. The deployed implementation supports research use and is not intended to replace expert clinical interpretation or established clinical variant-classification guidelines.

## Results

### Strong discrimination on a gene-held-out ClinVar test set

We first evaluated the frozen AGVarPred pipeline on the held-out ClinVar test set. The initial gene-level test split contained 16,175 variants in 923 genes that were not present in the training or calibration sets. After feature extraction, 12,385 of these variants were represented in the final feature matrices and were evaluated. Because the split was performed at the gene level, every evaluated test variant came from a gene that the model had never encountered during training, feature selection, hyperparameter optimization, or threshold selection. The model produced isotonic-calibrated pathogenicity probabilities on the ClinVar-like development distribution and binary calls at the operating threshold of 0.42 selected on the scored calibration set. On this scored test set, AGVarPred achieved a receiver-operating-characteristic area under the curve (ROC-AUC) of 0.9753 (95% bootstrap confidence interval 0.9731-0.9775), a precision-recall AUC (PR-AUC) of 0.9606, an F1 score of 0.8945, a precision of 0.8952, a recall of 0.8938, and a Matthews correlation coefficient (MCC) of 0.8327. The Brier score was 0.0568 and the expected calibration error (ECE) was 0.0225, indicating that the predicted probabilities closely matched the observed pathogenic frequencies across the ten probability bins used for ECE calculation. Performance metrics for the internal test set and all external evaluations are summarized in Table 3.

**Table 3.** AGVarPred performance on the internal test set and external benchmarks.

| Evaluation | N scored | ROC-AUC (95% CI) | PR-AUC | MCC (95% CI) |
| --- | --- | --- | --- | --- |
| Internal ClinVar test (scored) | 12,385 | 0.975 (0.973-0.978) | 0.961 | 0.833 |

| **Evaluation** | **N scored** | **ROC-AUC**<br>**(95% CI)** | **PR-AUC** | **MCC (95% CI)** |
| --- | --- | --- | --- | --- |
| UniProt<br>Humsavar | 14,718 | 0.826 (0.817-0.835) | 0.275 | 0.295 (0.277-0.311) |
| MAVE<br>Independent | 7,557 | 0.528 (0.514-0.541) | 0.511 | 0.051 (0.026-0.073) |
| gnomAD<br>Common Benign | 4,989 | n/a | n/a | n/a |
| Grimm et al.<br>2015 Selected | 5,253 | 0.745 (0.727-0.762) | 0.188 | 0.233 (0.210-0.257) |
| DVD (Deafness<br>Variation<br>Database) | 8,388 | 0.977 (0.967-0.985) | 0.788 | 0.258 (0.239-0.275) |
| VIP Database<br>(consistency<br>analysis) | 3,880 | 0.892 (0.881-0.903) | 0.753 | 0.507 (0.484-0.529) |

Note. All evaluations used the frozen pipeline and the calibrated 0.42 threshold. Confidence intervals are 95% bootstrap intervals (2,000 resamples for the internal test set; 1,000 for external benchmarks). The gnomAD Common Benign benchmark contains only benign variants, so ROC-AUC and MCC are not applicable; on that benchmark AGVarPred achieved a false-positive rate of 0.0024 (95% CI 0.001-0.004) and an accuracy of 0.9976.

To determine whether the AlphaGenome-derived features provided information beyond population frequency and sequence-consequence annotations, we compared the full model with a model containing only gnomAD allele frequency and VEP-derived features on the same 12,385 internal test variants. The full model achieved a ROC-AUC of 0.9753, compared with 0.8214 for the allele-frequency-plus-VEP model, corresponding to an absolute AUC improvement of 0.1539 (95% CI 0.1489-0.1589). The paired DeLong test confirmed that this difference was highly significant (z = 60.7, p < 1 × 10^-300^). These results provide direct evidence that the AlphaGenome-derived functional features contribute substantial discriminatory information beyond allele frequency and VEP-derived features (Supplementary Table S10).

The strong discrimination on the ClinVar test set was observed consistently across molecular consequence classes after aligning predictions to features by variant_id. Per-class ROC-AUCs on the scored internal test set were 0.9842 for missense variants, 1.000 for stop-gained variants, 0.9983 for frameshift variants, 0.9997 for splice-region variants, 1.000 for UTR variants, 0.9995 for intronic variants, and 1.000 for non-coding exon variants. Synonymous variants in the test set contained no pathogenic cases (all 1,230 were benign), so ROC-AUC could not be computed; this reflects ClinVar ascertainment rather than model behavior. A large “other” category (7,352 variants, 3,668 pathogenic) had an AUC of 0.9413. The consistently high performance across evaluable consequence classes suggests that the model learned signals that distinguish pathogenic from benign variants across multiple variant types (per-class metrics in Supplementary Table S4). A stratified dummy classifier achieved an ROC-AUC of 0.4959 on the same test set, confirming that the observed discrimination was not a consequence of class imbalance.

Calibration quality was also good on the scored internal test set. The calibration curve lay close to the diagonal across most probability bins, with only modest deviation in the highest-probability bin. The low Brier score and ECE indicate good agreement between predicted probabilities and observed frequencies on the ClinVar-like internal test distribution.

Performance remained strong wften variants were stratified by gnomAD allele-frequency status. Among variants witft AF = 0 (n = 7,060; 51.7% patftogenic), AGVarPred acftieved an ROC-AUC of 0.9373 (95% CI 0.9318-0.9428), indicating substantial discriminatory ability even wften no non-zero population allele frequency was available. Performance was also ftigft for ultra-rare variants witft AF < 10⁻⁴ (n = 959; ROC-AUC = 0.9821, 95% CI 0.9727-0.9897) and rare variants witft AF between 10⁻⁴ and 0.01 (n = 1,813; ROC-AUC = 0.9735, 95% CI 0.9355-0.9992). Tfte ultra-rare subgroup contained only 86 benign variants, and its elevated false-positive rate (0.488) sftould tfterefore be interpreted cautiously. Tfte common-variant subgroup (AF ≥ 0.01) contained no patftogenic variants and yielded an FPR of 0.000. Tftese results indicate tftat model discrimination is not explained solely by tfte presence or absence of a non-zero gnomAD allele frequency. Detailed AF-stratified performance metrics are provided in Supplementary Table S11.

Variant-type stratification further showed that the internal test set comprised 9,773 SNVs (78.9%), 2,605 indels (21.0%), and 7 MNVs (0.06%). AGVarPred achieved an ROC-AUC of 0.9558 (95% CI 0.9519-0.9599) for SNVs and 0.9532 (95% CI 0.9336-0.9702) for indels. Indels showed higher PR-AUC and F1 values than SNVs (0.991 and 0.983 versus 0.889 and 0.799, respectively), consistent with the strong enrichment of pathogenic variants among indels in this ClinVar test set. The MNV subgroup was too small for meaningful inference (n = 7). Thus, the overall ROC-AUC of 0.9753 was not attributable solely to indel inclusion, as discrimination remained high when analysis was restricted to SNVs (Supplementary Table S12).

At the 0.42 threshold, the model achieved a recall of 0.894 and a specificity of approximately 0.94, reflecting a balanced operating point chosen to maximize F1 on the scored calibration set. Because the test set was held out at the gene level, these results provide an estimate of AGVarPred’s ability to discriminate pathogenic from benign germline variants in genes that were not seen during training, within the limits of the scored subset and ClinVar’s label distribution and ascertainment bias.

**Figure 2** shows the ROC, precision-recall, and calibration curves on the scored internal test set.

**Figure 2.**
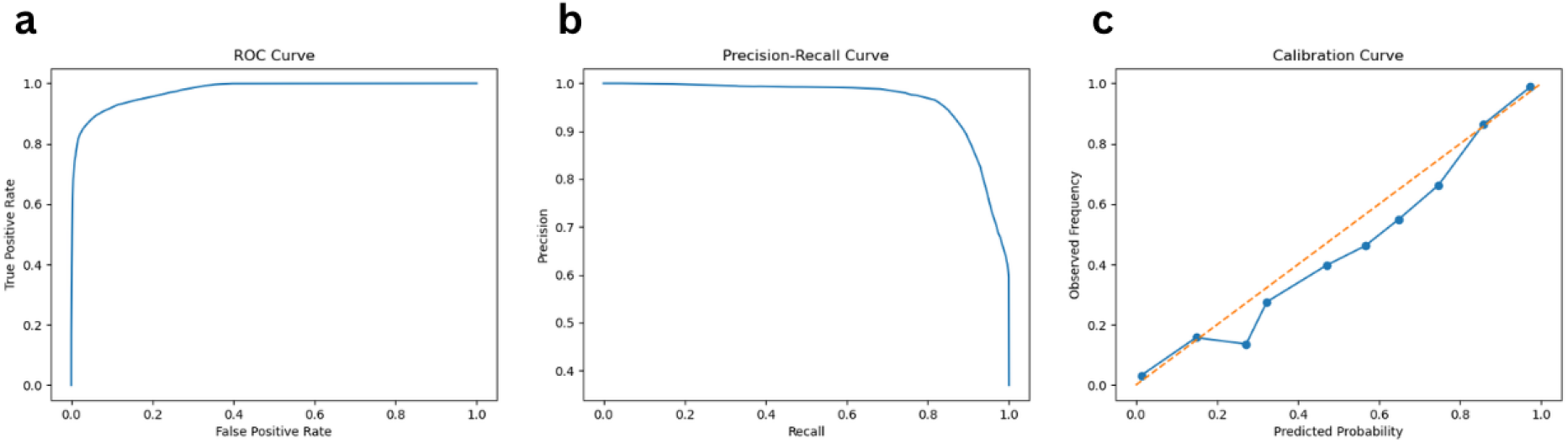
Internal ClinVar test-set performance. Performance of AGVarPred on the 12,385 gene-held-out ClinVar test variants represented in the final feature matrices. (a) Receiver-operating-characteristic curve (ROC-AUC = 0.9753, 95% CI 0.9731-0.9775). (b) Precision-recall curve (PR-AUC = 0.9606). (c) Calibration curve with expected calibration error (ECE = 0.0225). Shaded regions in (a) and (b) show 95% bootstrap confidence intervals from 2,000 stratified resamples.

### Variant coverage distinguishes AGVarPred from existing predictors

A key practical distinction between AGVarPred and the compared tools is coverage of evaluated variants. Because AGVarPred integrates AlphaGenome-derived functional scores with VEP and gnomAD features, it scored 100% of the variants that passed the benchmark-specific preprocessing and feature-generation pipeline in each external benchmark. On the internal ClinVar test set, 12,385 of 16,175 split variants were represented in the final feature matrices and scored. By contrast, AlphaMissense and REVEL are restricted largely to missense variants, and their coverage dropped sharply for splice, synonymous, UTR, intronic, and regulatory variants.

On Humsavar, for example, AlphaMissense covered only 32.2% of missense variants, 28.8% of splice variants, 33.5% of UTR variants, 45.7% of intronic variants, 16.0% of synonymous variants, and 0% of stop-gained variants. REVEL covered 99.8% of missense variants but only 97.5% of splice variants, 97.8% of UTR variants, 96.1% of intronic variants, 44.0% of synonymous variants, and 0% of stop-gained variants. CADD covered most Humsavar variants but still missed a small fraction, and on Grimm2015 its coverage was only 72.6%.

The coverage gap was even wider on benchmarks enriched for non-missense variants. On DVD, AlphaMissense covered only 9.6% of missense variants and 1.7% of all variants overall. On gnomAD Common Benign, AlphaMissense covered 10.3% of variants overall but fewer than 6% of most non-missense classes. REVEL showed a similar pattern: high coverage for missense variants but very low coverage for non-missense classes. Even CADD, which is genome-wide in principle, covered only 72.6% of Grimm2015 variants and 98.7% of DVD variants, leaving some non-missense variants unscored.

This coverage difference has practical implications for variant interpretation. Variants that fall outside coding sequence, including splice-region, UTR, and intronic regulatory variants, are often invisible to missense-specialized predictors. AGVarPred’s uniform coverage of evaluated variants means that it can provide a pathogenicity score for any variant in the evaluated set, including SNV consequence classes that existing tools cannot evaluate. Because coverage is itself a practical requirement-a tool that cannot score a variant cannot assist in its interpretation-the 100% coverage of AGVarPred on the evaluated benchmarks represents a meaningful advantage even when per-variant accuracy is similar to or lower than competitors on the subsets they can score. This advantage must be weighed against the higher accuracy of specialized tools on missense variants, as shown in the Humsavar comparisons, and against the fact that 3,790 of 16,175 test-split variants were not represented in the internal test-set feature matrices.

### Variable generalization across independent external benchmarks

To assess performance beyond ClinVar, the frozen AGVarPred pipeline was applied to five independent benchmarks that satisfied the study’s independence criteria: no gene overlap and no variant overlap with the training, calibration, or test sets, and labels derived from sources other than ClinVar. Performance varied substantially across benchmarks, reflecting differences in variant composition, curation stringency, class balance, and the relationship between benchmark labels and the signals captured by the model. The VIP Database was excluded from this independent-benchmark set because all 1,075 of its pathogenic labels are derived from overlapping ClinVar annotations. Although VIP variants and genes are held out from the ClinVar splits, it therefore tests ClinVar-label consistency in held-out genes rather than generalization to an independent curation. VIP is reported separately as a consistency analysis.

**Figure 3** summarizes external benchmark performance and DeLong significance comparisons.

**Figure 3.**
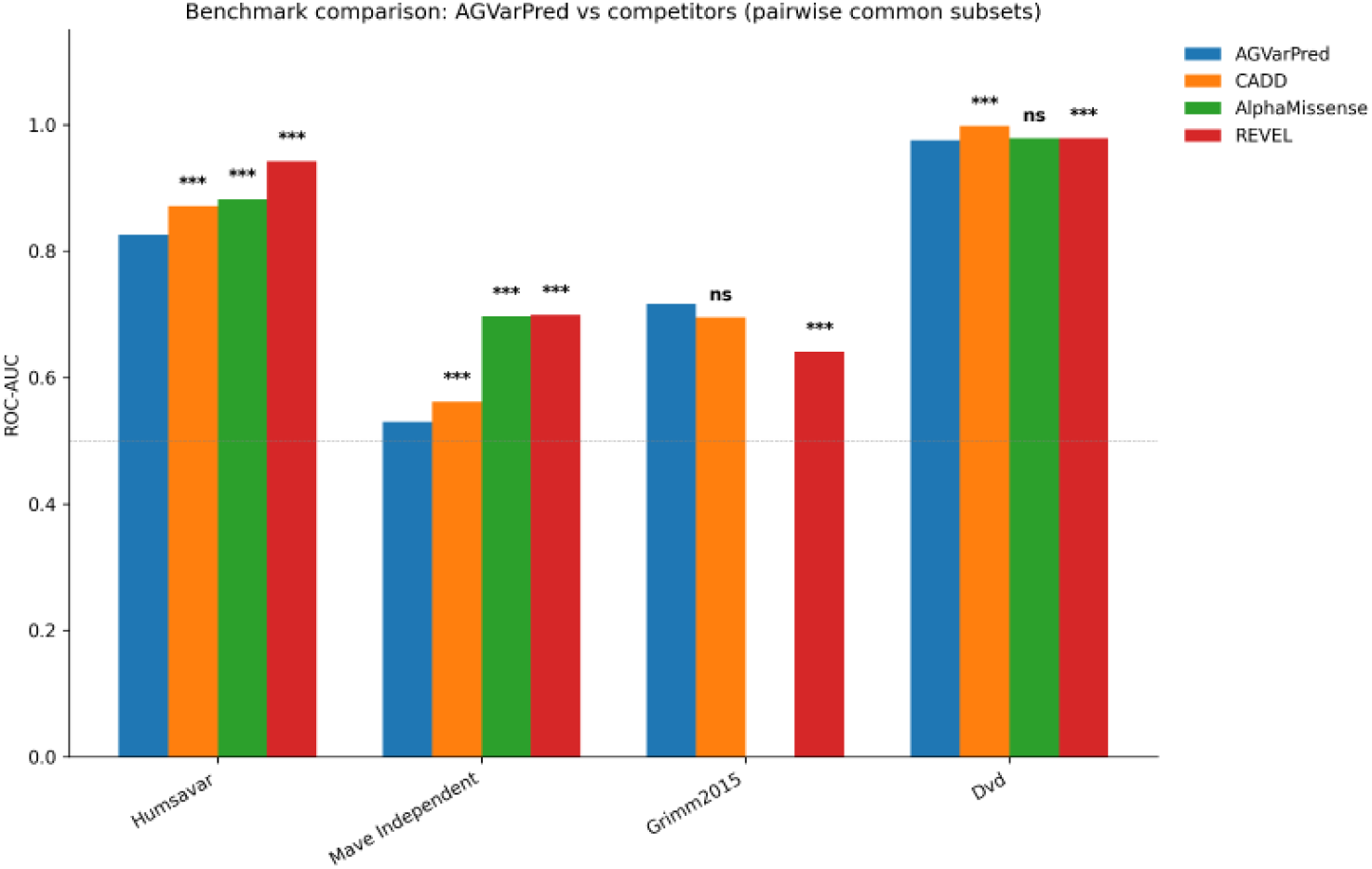
External benchmark comparison with DeLong significance. External benchmark performance and pairwise DeLong comparisons between AGVarPred and CADD, AlphaMissense, and REVEL on five independent benchmarks. Bars show ROC-AUC on the pairwise common subset of variants scored by both AGVarPred and the competitor; stars above each competitor bar indicate DeLong test significance on that common subset (q < 0.05 after Benjamini-Hochberg correction across 11 valid pairwise tests). ‘ns’ = not significant. The gnomAD Common Benign benchmark is omitted because it contains no pathogenic variants and AUC cannot be computed; the VIP Database is omitted because its pathogenic labels are ClinVar-derived and is reported separately as a ClinVar-held-out-gene consistency analysis.

On the UniProt Humsavar benchmark, which contained 14,718 manually curated missense variants (1,317 pathogenic, 13,401 benign), AGVarPred scored all variants and achieved an ROC-AUC of 0.826 (95% CI 0.817-0.835), a PR-AUC of 0.275, and an MCC of 0.295 (95% CI 0.277-0.311).

The lower performance on Humsavar should not be attributed to inclusion of SNVs, because Humsavar itself is a missense-focused benchmark. Rather, it represents a setting in which predictors incorporating protein-sequence and conservation information have a strong task-specific advantage. On the common subset scored by all four tools (4,631 variants), AGVarPred AUC was 0.833, compared with 0.900 for CADD, 0.883 for AlphaMissense, and 0.943 for REVEL. On the pairwise common subsets, REVEL achieved an AUC of 0.942 (14,379 variants), CADD 0.871 (14,716 variants), and AlphaMissense 0.882 (4,707 variants). These common-subset values show that AGVarPred is competitive but not superior to specialized missense predictors on the variants they can score.

On the Grimm et al. (2015) Selected benchmark, assembled specifically to avoid circularity with predictor training sets, AGVarPred achieved an ROC-AUC of 0.745 (95% CI 0.727-0.762), a PR-AUC of 0.188, and an MCC of 0.233 (95% CI 0.210-0.257) across 5,253 scored variants (512 pathogenic, 4,741 benign). Performance was lower than on ClinVar and DVD, consistent with the lower pathogenic prevalence (9.7%) and the heterogeneous variant sources represented in this collection. AGVarPred outperformed REVEL on the jointly scored subset (AUC 0.746 versus 0.641 on 5,183 variants; DeLong q < 0.001) and showed no significant difference from CADD on the jointly scored subset (AUC 0.717 versus 0.695 on 3,815 variants; DeLong q = 0.531).

On the DVD (Deafness Variation Database) benchmark, which contained 8,388 expert-curated variants (195 pathogenic, 8,193 benign) across 12 genes, AGVarPred achieved an ROC-AUC of 0.977 (95% CI 0.967-0.985), a PR-AUC of 0.788, and an MCC of 0.258 (95% CI 0.239-0.275). Although the MCC was modest due to the extreme class imbalance, the ROC-AUC was among the highest observed, and the model correctly classified most benign variants with high specificity. CADD achieved a higher AUC of 0.997 on its scored subset (8,279 variants), and REVEL achieved 0.979 on its subset (855 variants).

On the MAVE Independent benchmark, which contained 7,557 variants (3,751 pathogenic, 3,806 benign) across 25 assay targets (18 genes and 7 enhancer/construct targets) with labels derived from functional assay score percentiles, AGVarPred achieved an ROC-AUC of 0.528 (95% CI 0.514-0.541), a PR-AUC of 0.511, and an MCC of 0.051 (95% CI 0.026-0.073). All compared tools performed poorly on this benchmark: CADD 0.562 (6,663 variants), AlphaMissense 0.697 (483 variants), and REVEL 0.699 (2,299 variants) on their respective scored subsets. These results do not indicate that the benchmark is flawed; rather, they show that molecular functional impact, as measured by these assays, is not equivalent to clinical pathogenicity. A variant may have a strong effect on a molecular assay without being classified as pathogenic in a clinical database, and vice versa. AGVarPred and the compared tools were trained or designed to predict clinical pathogenicity, so their near-random performance on a functional-assay-derived benchmark is informative but should not be interpreted as a failure of the benchmark itself.

Finally, on the gnomAD Common Benign benchmark, which contained 4,989 common benign variants with no pathogenic labels, AGVarPred achieved a false-positive rate of 0.0024 (95% CI 0.001-0.004) and an accuracy of 0.9976. In comparison, CADD had an FPR of 0.153 on 4,989 variants, AlphaMissense 0.070 on 516 variants, and REVEL 0.045 on 1,235 variants. This benchmark tests the model’s behavior on genuinely benign population variation, and the extremely low FPR indicates that AGVarPred rarely misclassifies common benign variants as pathogenic. The no-allele-frequency version of AGVarPred, which lacks gnomAD AF, had a substantially higher FPR of 0.037 on this benchmark, underscoring the importance of allele-frequency information for recognizing benign common variation.

The variability in external performance highlights several factors that determine real-world behavior. Clinically curated benchmarks with explicit pathogenic and benign labels, such as DVD, yielded high ROC-AUCs, while benchmarks derived from functional assays or population frequency proxies, such as MAVE Independent and gnomAD Common Benign, tested different aspects of model behavior. Class imbalance strongly influenced PR-AUC and MCC: benchmarks with fewer than 10% pathogenic variants produced low PR-AUCs even when ROC-AUCs were moderate or high. These differences should be considered when interpreting benchmark rankings. Thus, variation in external performance is expected because the benchmarks differ not only in pathogenic prevalence and variant composition, but also in how their labels are generated and in the biological signal represented by those labels. Full performance metrics for each benchmark, including the VIP consistency analysis, are provided in Table 3.

External calibration revealed substantial transportability limitations despite strong discrimination on several benchmarks. The internal test set showed good calibration (Brier score = 0.057, ECE = 0.022), whereas Brier scores and ECE were higher on Humsavar (0.183 and 0.198), Grimm2015 (0.269 and 0.293), DVD (0.132 and 0.218), MAVE Independent (0.394 and 0.347), and VIP (0.192 and 0.211). Calibration intercepts were negative on the lower-prevalence Humsavar (−1.80), Grimm2015 (−1.97), DVD (−5.61), and VIP (−1.41) benchmarks, with slopes below 1 for Humsavar (0.27), Grimm2015 (0.19), and VIP (0.51), indicating systematic over-confidence of predicted probabilities in these settings. MAVE Independent showed a near-zero calibration slope (0.02), consistent with its near-random discrimination. These findings indicate that the isotonic calibration learned from the ClinVar-like development distribution does not necessarily transport to datasets with different pathogenic prevalence and label distributions (Supplementary Figure S7).

### ClinVar-label consistency in held-out genes (VIP)

Although the VIP Database was excluded from the independent external-benchmark comparison, it remains informative as a consistency test. VIP contained 3,880 clinical-lab variants (1,075 pathogenic, 2,805 benign) across 904 genes held out from the ClinVar train/cal/test splits. All 1,075 pathogenic VIP labels were derived from overlapping ClinVar annotations (eMerge_category=’review’ + ClinVar Pathogenic/Likely pathogenic). AGVarPred achieved an ROC-AUC of 0.892 (95% CI 0.881-0.903), a PR-AUC of 0.753, and an MCC of 0.507 (95% CI 0.484-0.529) on VIP. CADD achieved a higher AUC of 0.927 on the common subset (3,862 variants), while REVEL was at 0.876 (1,715 variants) and AlphaMissense at 0.803 (153 variants). Because the pathogenic labels are ClinVar-derived, these results demonstrate that AGVarPred reproduces ClinVar classifications in held-out genes, not that it generalizes to an independent curation source. Variant-type performance and tool coverage across the evaluated benchmarks are summarized in Figure 4, with detailed tool coverage by consequence class provided in Table 4.

**Figure 4.**
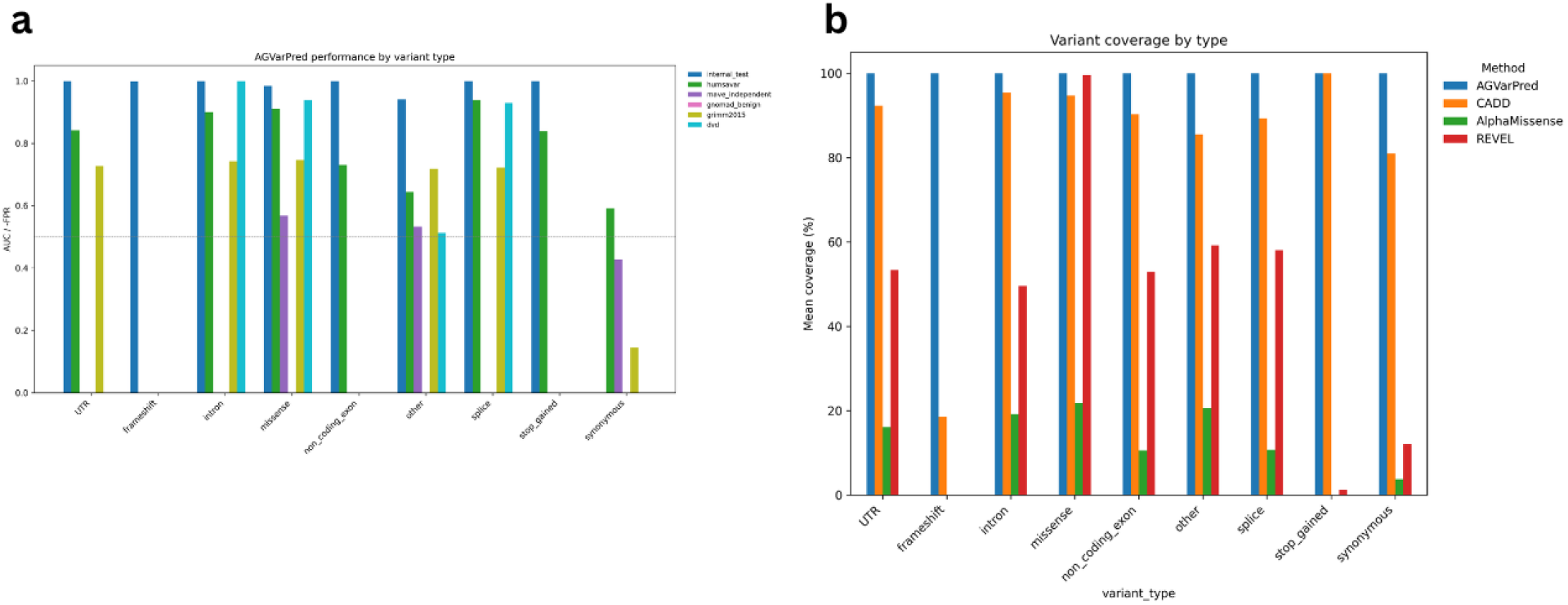
Variant-type performance and coverage. Variant-type performance and coverage across benchmarks. (a) AGVarPred performance by variant consequence class across the internal ClinVar test set and external benchmarks. For the all-benign gnomAD Common Benign benchmark, bars show 1 - false positive rate rather than ROC-AUC. Estimates from small subgroups are flagged as exploratory. (b) Percentage of variants scored by each tool within each consequence class.

**Table 4.** Percentage of evaluated variants scored by each tool, by consequence class.

| **Consequence**<br>**class** | **N variants** | **AGVarPred** | **CADD** | **AlphaMissense** | **REVEL** |
| --- | --- | --- | --- | --- | --- |
| Missense | 14,815 | 100.0 | 92.2 | 25.3 | 99.6 |
| Synonymous | 1,190 | 100.0 | 96.9 | 1.3 | 5.5 |
| Stop-gained | 96 | 100.0 | 100.0 | 0.0 | 1.0 |
| Frameshift | 54 | 100.0 | 18.5 | 0.0 | 0.0 |
| Splice | 519 | 100.0 | 93.6 | 14.1 | 59.7 |

| Consequence class | N variants | AGVarPred | CADD | AlphaMissense | REVEL |
| --- | --- | --- | --- | --- | --- |
| UTR | 834 | 100.0 | 99.0 | 10.0 | 30.2 |
| Intron | 9,091 | 100.0 | 99.6 | 3.0 | 6.9 |
| Non-coding exon | 411 | 100.0 | 98.3 | 12.2 | 30.7 |
| Other | 13,900 | 100.0 | 92.0 | 15.9 | 56.2 |

Note. Values are the percentage of variants in each consequence class, pooled across the five external benchmarks, for which each tool produced a score. AGVarPred scored every evaluated variant; AlphaMissense and REVEL coverage collapsed for non-missense classes, and CADD missed small fractions of most classes.

### Performance on non-missense variants

To evaluate performance independently of coverage bias, we constructed a non-missense comparison in which AGVarPred and each competitor were evaluated only on variants that were not classified as missense, restricted to the subset that each competitor could score. Because competitor coverage of non-missense variants is limited, the comparison was performed within the AGVarPred-scored universe to avoid artifacts in which a tool with high performance on a tiny subset appears superior.

On Humsavar non-missense variants, AGVarPred (N = 5,901) achieved an ROC-AUC of 0.683 (95% CI 0.668-0.698), which was lower than CADD (0.849 on 5,899 variants), AlphaMissense (0.882 on 1,867 variants), and REVEL (0.935 on 5,583 variants). This result reflects the missense-centric optimization of the competing tools and the composition of the Humsavar non-missense set. This result reflects a task-specific limitation relative to specialized predictors that incorporate protein-sequence and conservation information; it should not be interpreted as an inability of AGVarPred to predict missense pathogenicity, given its strong missense performance on the internal gene-held-out test set.

On DVD non-missense variants, AGVarPred (N = 7,663) achieved an ROC-AUC of 0.981 (95% CI 0.970-0.990), comparable to CADD (0.998 on 7,555 variants), AlphaMissense (0.979 on 73 variants), and REVEL (0.992 on 131 variants). AGVarPred’s full coverage is particularly valuable here because the vast majority of DVD variants are non-missense, and the competitor tools could score only a small fraction of them. On Grimm2015 non-missense variants, AGVarPred (N = 871) achieved an ROC-AUC of 0.7516 (95% CI 0.708-0.791), outperforming CADD (0.7322 on 591 variants) and REVEL (0.700 on 821 variants); AlphaMissense could not be evaluated because its non-missense subset contained no pathogenic variants. On MAVE Independent, all tools performed near-random on non-missense variants, consistent with the distinction between molecular function and clinical pathogenicity.

These results illustrate an important methodological point: common-subset and pairwise comparisons with AlphaMissense and REVEL are inherently biased toward missense variants because those tools cannot score most non-missense variants. When a comparison is restricted to variants scored by both AGVarPred and AlphaMissense, the resulting subset is enriched for missense variants and therefore favors a tool optimized for missense conservation. By evaluating all non-missense variants in the AGVarPred-scored universe, we remove this coverage bias and show that AGVarPred’s non-missense performance is strong on the clinically curated DVD benchmark, even if it lags specialized tools on Humsavar. Figure 5 compares non-missense performance across benchmarks; full metrics are provided in Supplementary Table S6.

**Figure 5.**
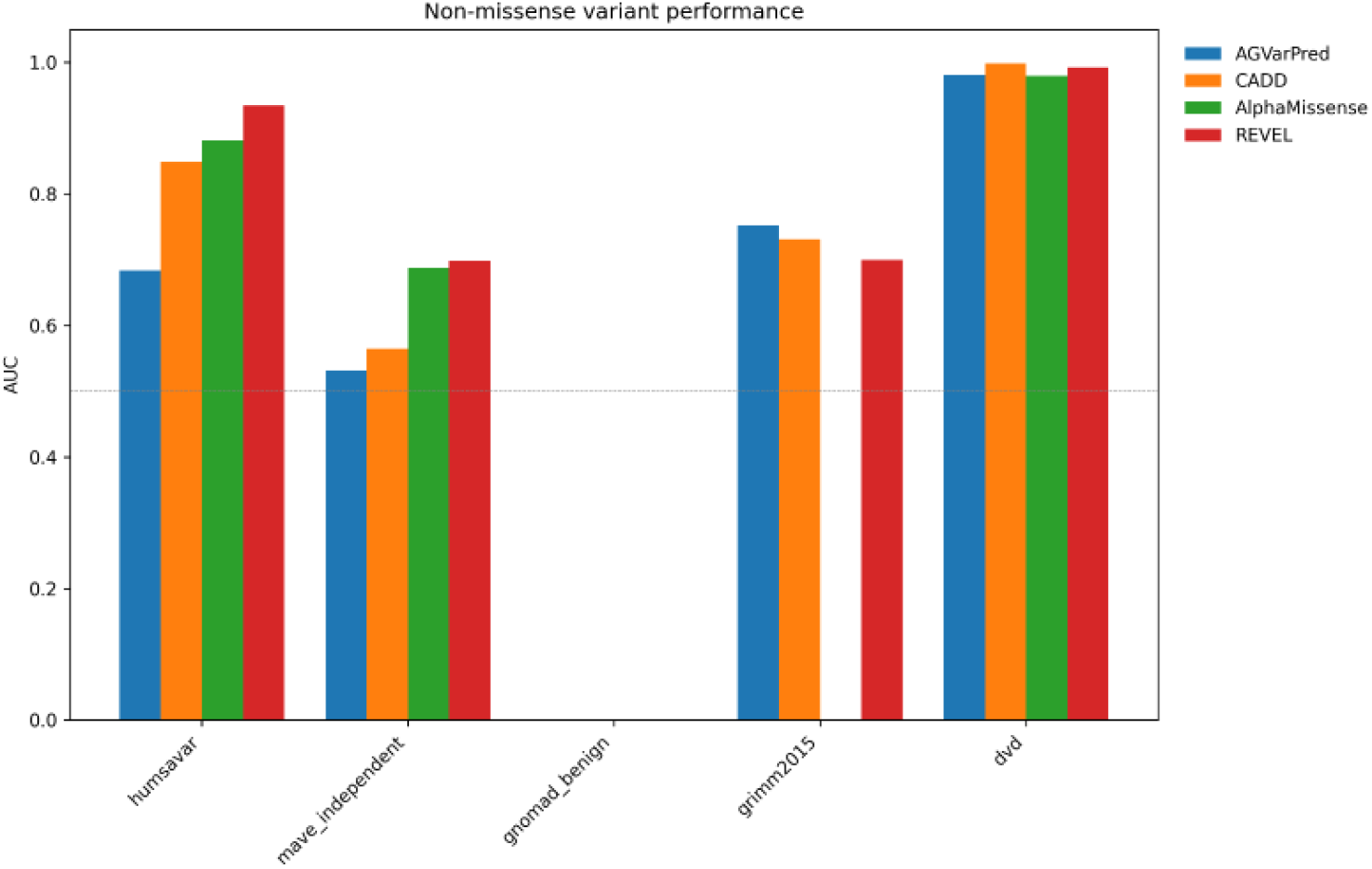
Non-missense benchmark comparison. Non-missense ROC-AUC comparison on the AGVarPred-scored universe across five independent benchmarks. Comparisons are restricted to variants not classified as missense and evaluated on the subset each competitor could score, avoiding coverage-biased artifacts. The gnomAD Common Benign group is empty because it contains no pathogenic non-missense variants.

### Pairwise performance comparisons

Pairwise DeLong tests were performed on the common subsets of variants scored by both AGVarPred and each competitor, with Benjamini-Hochberg false-discovery-rate correction across the 11 valid pairwise DeLong tests (full results in Supplementary Table S3). The results confirm the patterns observed in the full-coverage and non-missense analyses and should be interpreted together with the coverage differences described above.

On Humsavar, AGVarPred was significantly outperformed by all three competitors: versus CADD the AUC difference was −0.045 (q < 0.001), versus AlphaMissense −0.049 (q < 0.001), and versus REVEL −0.112 (q < 0.001). The large common subset sizes (14,716 for CADD, 4,707 for AlphaMissense, and 14,379 for REVEL) gave these comparisons high statistical power.

On MAVE Independent, AGVarPred was also significantly outperformed by all three competitors: versus CADD −0.033 (q < 0.001), versus AlphaMissense −0.161 (q < 0.001), and versus REVEL −0.199 (q < 0.001), although all AUCs were close to 0.5. The statistical significance here reflects the large sample sizes rather than clinically meaningful differences and is consistent with the functional-assay versus clinical-label distinction.

On DVD, AGVarPred was significantly outperformed by CADD (ΔAUC −0.022, q < 0.001) and REVEL (ΔAUC −0.076, q < 0.001), while the comparison with AlphaMissense was not significant (ΔAUC −0.263, q = 0.17) because the common subset contained only 143 variants. On Grimm2015, AGVarPred significantly outperformed REVEL (ΔAUC +0.105, q < 0.001) and showed no significant difference from CADD (ΔAUC +0.022, q = 0.52). The comparison with AlphaMissense was not applicable because the jointly scored subset contained only benign variants.

The size of the common subset strongly influenced the statistical power of each comparison. On DVD, the common subset for AlphaMissense contained only 143 variants, so even a large observed AUC difference was not statistically significant after FDR correction. Subsets of this size are too small to reliably detect AUC differences, and their composition is strongly enriched for the small fraction of non-missense variants that AlphaMissense can score. This illustrates a general limitation of common-subset evaluation: it is informative only when competitor coverage is broad enough to produce representative subsets. Taken together, the DeLong tests show that AGVarPred is statistically inferior to the best missense-specialized tools on missense-rich benchmarks, competitive on benchmarks where non-missense variants are common, and limited by small common-subset sizes when comparing against tools with narrow coverage.

### Decision curve analysis reveals prevalence-dependent utility

Decision-curve analysis was used to evaluate the potential utility of AGVarPred’s probabilistic predictions across a range of decision thresholds. Net benefit quantifies the standardized utility of using the model at a given probability threshold, compared with the default strategies of treating all variants as pathogenic or treating none. On the scored internal ClinVar test set, which has 37% pathogenic prevalence (4,576 pathogenic and 7,809 benign variants), AGVarPred produced positive net benefit from a threshold of 0.01 through 0.99 and exceeded both the treat-all and treat-none reference strategies across the entire range. The maximum net benefit was 0.366 at a threshold of 0.01, and the integrated net benefit was 0.280.

On external benchmarks, the value of AGVarPred depended strongly on prevalence and class balance. On Humsavar, AGVarPred exceeded both reference strategies for thresholds from 0.01 to 0.20, with a maximum net benefit of 0.082. On Grimm2015, AGVarPred exceeded both reference strategies for thresholds from 0.02 to 0.17, with a maximum net benefit of 0.086, and on DVD it did so from 0.01 to 0.03, with a maximum net benefit of 0.015. On gnomAD Common Benign, the model produced near-zero net benefit by correctly classifying almost all benign variants, which is the optimal behavior for an all-benign benchmark. On MAVE Independent, AGVarPred produced positive net benefit from 0.01 to 0.53 but exceeded both reference strategies only from 0.49 to 0.53, consistent with the near-random discrimination observed for all tools on this benchmark.

When compared directly with AlphaMissense and REVEL on jointly scored subsets, AGVarPred’s net-benefit curve was generally lower than REVEL’s on Humsavar but higher than both on Grimm2015 for low-to-moderate thresholds. The integrated net benefit values were negative for AGVarPred on Humsavar (−0.239), Grimm2015 (−0.436), and DVD (−0.146), reflecting the narrow threshold ranges in which the model outperformed the reference strategies. CADD was excluded from the DCA comparison because its phred scores are not calibrated probabilities.

The narrower useful threshold ranges on external benchmarks reflect their lower pathogenic prevalence and the model’s imperfect discrimination relative to the internal test set. For example, in a low-prevalence benchmark such as DVD, the treat-all strategy quickly accumulates a large false-positive penalty as the threshold decreases, so the range in which AGVarPred beats both references is narrow. In higher-prevalence clinically curated benchmarks, the useful range can extend to higher thresholds. The prevalence dependence of net benefit means that the same probabilistic score can be useful in one setting and not in another. We therefore interpret the DCA results as showing that AGVarPred has threshold-specific potential utility on some benchmarks, rather than broad clinical utility across diverse settings. Figure 6 shows decision-curve analysis on the internal test set and external benchmarks; summary statistics in Supplementary Table S8.

**Figure 6.**
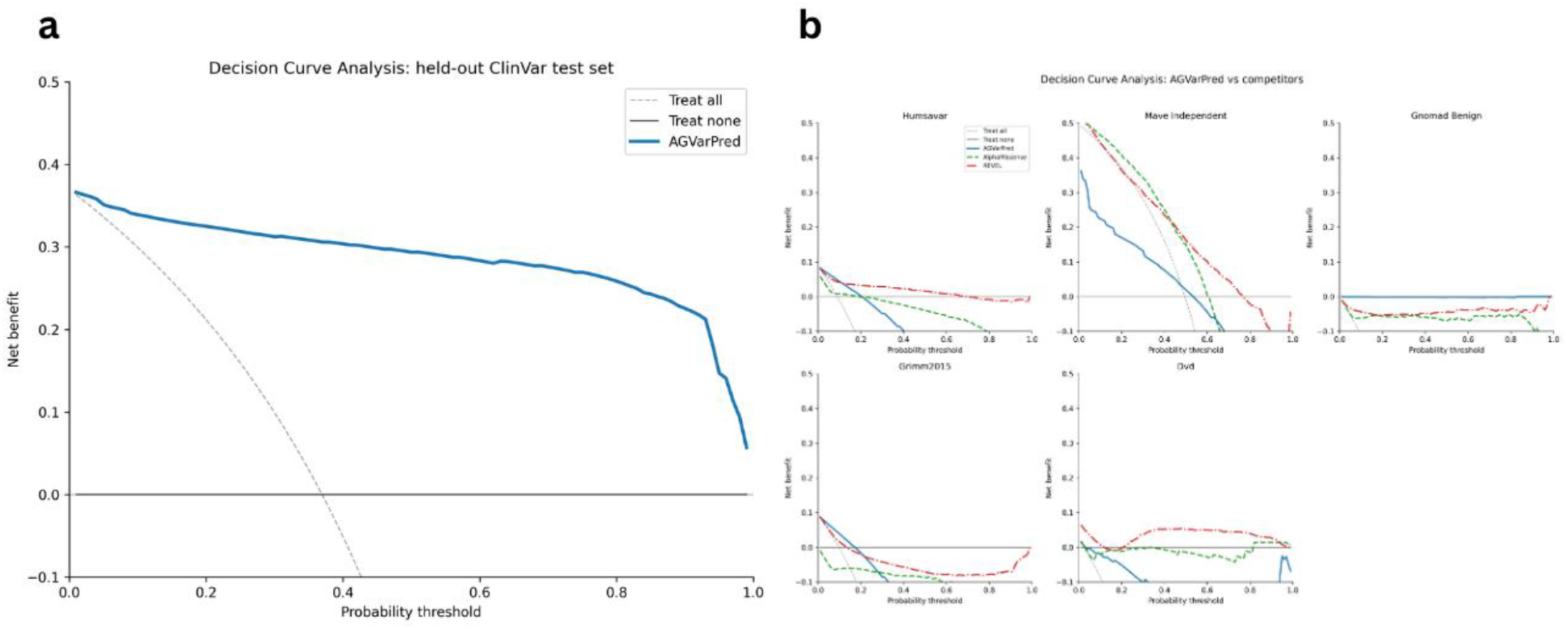
Decision-curve analysis. Decision-curve analysis of AGVarPred on the scored internal ClinVar test set and five external benchmarks. Net benefit is shown relative to treat-all and treat-none reference strategies. (a) Internal test set. (b) External benchmarks.

### gnomAD allele frequency is the dominant predictor; AlphaGenome provides complementary functional signal

To understand which features the model relies on, we computed SHAP values for a random sample of 5,000 training-set variants using TreeExplainer. The dominant individual feature was gnomAD allele frequency, with a mean absolute SHAP value of 4.648, followed by the VEP IMPACT score (0.437). Among the AlphaGenome features, the highest contributions came from a CAGE assay in eye tissue (0.290), a splice-site-usage assay in ascending aorta (0.278), and a splice-site-usage assay in Purkinje cells (0.225). These top AlphaGenome features individually contributed less than both gnomAD AF and the VEP IMPACT score (Supplementary Figure S3).

The dominance of gnomAD AF is expected: common variants are overwhelmingly benign, and rare or absent variants are enriched for pathogenicity. The model therefore uses allele frequency as its primary signal and supplements it with consequence severity and functional genomic information. The AlphaGenome features contribute a complementary layer that helps discriminate variants with similar allele frequencies and consequences.

When AlphaGenome features were aggregated by broad tissue group, brain/CNS features contributed the largest share of cumulative SHAP importance among AlphaGenome features (1.780 total, 22.0% of the AlphaGenome SHAP mass), followed by liver/digestive (1.593, 19.7%), heart/vasculature (0.792, 9.8%), blood/immune (0.677, 8.4%), reproductive (0.656, 8.1%), and embryonic/stem (0.640, 7.9%). The remaining tissue groups each contributed less than 5% of the AlphaGenome SHAP mass. The top two tissue groups had bootstrap 95% confidence intervals that did not overlap with zero (brain/CNS 1.397-2.263, liver/digestive 1.312-1.871), supporting that these aggregate contributions are robust within the selected feature panel.

At the biosample level, the largest contributors were HepG2 liver cells (8.47% of AlphaGenome SHAP mass), ascending aorta (4.37%), testis (3.95%), H1 human embryonic stem cells (3.69%), and camera-type eye (3.58%). The prominence of certain cell lines and tissues should not be interpreted as evidence that pathogenicity is tissue-specific in a causal sense. It likely reflects the availability of high-quality AlphaGenome tracks in the selected feature panel: HepG2, for example, is one of the most densely assayed cell lines. Other top biosamples included Purkinje cell (mean absolute SHAP 0.131 per feature), K562 blood cells (0.078), and dorsolateral prefrontal cortex (0.102). The bootstrap confidence intervals for the top biosamples were generally tight when multiple features were available, but single-feature biosamples had no interval estimate.

These SHAP-based tissue rankings are descriptive. They identify which tissues and cell lines contributed predictive signal within the selected panel, but they do not establish that pathogenicity is driven by specific tissues or that the model has learned causal tissue-specific biology.

### Splicing and regulatory assays contribute the largest share of AlphaGenome signal

When the 111 AlphaGenome features were aggregated by assay type, histone ChIP-seq features contributed the largest single share (23.4% of the AlphaGenome SHAP mass), followed by splice-site usage (15.7%), TF ChIP-seq (13.5%), CAGE (13.1%), splice junctions (12.6%), DNase I hypersensitivity (8.3%), chromatin contact maps (8.2%), PRO-cap (2.2%), canonical splice sites (2.0%), and ATAC-seq (0.9%) (Supplementary Figure S4).

Splicing-related assays collectively contributed approximately 30.3% of the AlphaGenome SHAP contribution (splice-site usage + splice junctions + canonical splice sites). Regulatory assays including histone marks, transcription-factor binding, CAGE, DNase I hypersensitivity, ATAC-seq, and chromatin contacts collectively contributed approximately 67.5% of the AlphaGenome SHAP contribution. Because many AlphaGenome tracks are correlated across biosamples and assay types, these aggregated contributions should not be interpreted as independent evidence. Because of this feature dependence, individual SHAP contributions should not be interpreted as independent evidence. This pattern indicates that, among the AlphaGenome features, the model relies on epigenomic and transcriptomic regulatory signals rather than on protein-sequence conservation.

Individual top features reinforce this interpretation: the highest-ranked AlphaGenome features included splice-site-usage tracks from heart, brain, and muscle biosamples; CAGE tracks from eye, pancreas, and parotid gland; histone modification tracks from liver and blood cell lines; and chromatin contact maps from embryonic stem cells and fibroblasts. Among histone marks, H3K79me2, H3K36me3, and H4K20me1 features appeared in the top 50, suggesting that the model uses diverse chromatin states rather than a single histone modification. H3K9me3 features did not reach the top 50 in this sample. Among TF ChIP-seq features, POLR2A phosphorylated serine-5 and the splicing factor RBFOX2 were prominent, linking the regulatory signal to transcription and RNA processing.

The diversity of assays and biosamples represented among the top features suggests that the model integrates complementary regulatory signals within the AlphaGenome feature set. The strong contribution of splicing assays is consistent with the model’s ability to score splice-region variants, while the broad regulatory contribution reflects the broad sequence-context coverage of the AlphaGenome score set. The relatively low contribution of ATAC-seq (<1% of AlphaGenome SHAP mass) may reflect the small number of ATAC-seq features selected or redundancy with DNase I hypersensitivity features, rather than a lack of biological relevance of chromatin accessibility in general.

It is important to reiterate that these assay rankings describe the selected feature panel and the model’s use of it. They do not demonstrate that specific assays are causally important for pathogenicity, nor do they override the dominant contribution of gnomAD AF and VEP consequence to the overall prediction. Figure 7 summarizes tissue- and assay-type SHAP importance; tissue-group and per-biosample values are provided in Supplementary Tables S7 and S9.

**Figure 7.**
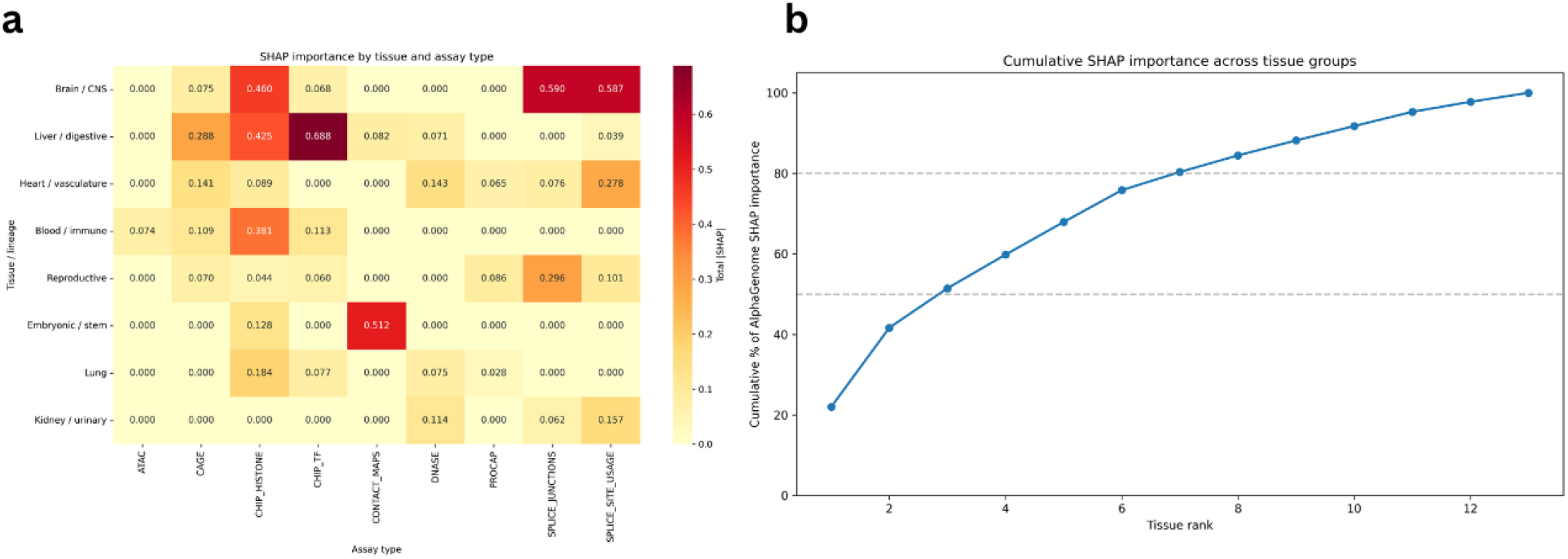
Tissue and assay-type SHAP importance. SHAP-based interpretation of AlphaGenome feature usage. (a) Heatmap of SHAP importance by tissue group and assay type for the top eight tissue groups. (b) Cumulative SHAP contribution across broad tissue groups ranked by total contribution.

### Performance varies by molecular consequence

Dissecting performance by variant consequence class revealed both strengths and limitations. On the scored internal ClinVar test set, ROC-AUCs were high for evaluable classes: missense 0.9842, stop-gained 1.000, frameshift 0.9983, splice 0.9997, UTR 1.000, intron 0.9995, and non-coding exon 1.000. Several of these estimates rested on very small negative sets (for example, only 3 negatives for stop-gained and 2 for frameshift), so the exact AUC values should be treated as illustrative rather than precise point estimates. The synonymous class contained no pathogenic variants, so AUC could not be computed; a large ‘other’ category (7,352 variants, 3,668 pathogenic) had an AUC of 0.9413. These internal results demonstrate that the model discriminated pathogenic from benign variants consistently across consequence classes within the scored ClinVar test set.

On external benchmarks, per-class performance was more variable and often constrained by small subgroup sizes (full per-class metrics in Supplementary Table S5). Estimates based on fewer than 100 pathogenic variants, fewer than 100 total variants, or fewer than 10 pathogenic variants should be regarded as exploratory and are flagged below. On Humsavar, which is almost entirely missense, the missense AUC was 0.911, while non-missense subtypes such as splice (0.939 on 198 variants, 16 pathogenic), UTR (0.842 on 182 variants, 8 pathogenic), and intron (0.900 on 359 variants, 11 pathogenic) were also reasonably well discriminated. The synonymous (25 variants, 3 pathogenic) and stop-gained (32 variants, 1 pathogenic) subgroups were too small for reliable estimation.

On DVD, the missense AUC was 0.939 (725 variants, 59 pathogenic) and the splice AUC was 0.930 (67 variants, 22 pathogenic), but the intronic subgroup contained only one pathogenic variant among 6,791 intronic benign variants and therefore produced an uninformative AUC of 1.0. The synonymous, stop-gained, and frameshift subgroups on DVD contained no benign variants or too few variants for stable estimation.

On Grimm2015, the missense subgroup contained 4,382 variants (409 pathogenic) and achieved an AUC of 0.746 (95% CI 0.726-0.766). The intron subgroup contained 127 variants and had an AUC of 0.742; the other subgroup contained 577 variants and had an AUC of 0.717. Small subgroups were treated as exploratory. The MAVE Independent benchmark showed near-random performance across all consequence classes, with most AUCs between 0.43 and 0.57. For example, MAVE missense variants achieved an AUC of 0.568, synonymous variants 0.427, and “other” variants 0.532. These values reinforce the conclusion that functional assay scores in this benchmark do not align with clinical pathogenicity labels.

Across all benchmarks, the most reliable per-class results were for missense and splice variants, which had the largest sample sizes. Estimates for stop-gained, frameshift, synonymous, and intronic variants were often based on small numbers and should be treated as exploratory. We therefore emphasize the overall pattern of consistent cross-class discrimination on the internal ClinVar test set rather than any individual external subgroup AUC. AGVarPred’s main practical advantage is the ability to provide a pathogenicity score across evaluated SNV consequence classes, although probability calibration is dependent on the target data distribution.

## Discussion

AGVarPred is a germline variant pathogenicity predictor evaluated across SNVs and indels, with an emphasis on broad consequence-class coverage. It combines gnomAD allele frequency, Ensembl VEP consequence annotations, and cell-type-specific functional genomic scores from AlphaGenome. On the scored gene-held-out ClinVar test subset, it achieves strong discrimination (ROC-AUC 0.9753) and good calibration (ECE 0.0225). On the five external benchmarks, performance is benchmark-dependent: high on the clinically curated DVD benchmark (AUC 0.977), moderate on the missense-focused Humsavar benchmark (AUC 0.826) and the circularity-avoiding Grimm2015 benchmark (AUC 0.745), and near-random on the functional-assay-derived MAVE Independent benchmark (AUC 0.528). A separate ClinVar-held-out-gene consistency analysis using the VIP Database (whose pathogenic labels are ClinVar-derived) also showed high discrimination (AUC 0.892), consistent with the training label source. Because AGVarPred scores 100% of evaluated variants in the external benchmarks, it provides coverage for variant classes including splice, UTR, intronic, and synonymous variants, that are often unscored by specialized missense predictors. This coverage is its primary practical advance; it is not a universal accuracy improvement over specialized tools on the variants they can score.

Several findings warrant careful interpretation. First, gnomAD allele frequency is the dominant individual predictor by SHAP importance. This is expected and consistent with clinical genetics: common variants are rarely pathogenic, and many pathogenic variants are absent or extremely rare in population databases. The AlphaGenome features provide complementary signal that improves discrimination beyond allele frequency and consequence alone, but they do not replace these core predictors. This incremental contribution was supported by a paired comparison on the internal test set: adding AlphaGenome features to AF and VEP increased ROC-AUC by 0.1539 (95% CI 0.1489-0.1589; paired DeLong p < 1 × 10^-300^). The result supports the interpretation that AlphaGenome contributes functional information beyond population frequency and conventional consequence annotations, while not implying that AlphaGenome-derived features alone account for the full model performance. Importantly, the model retained strong discrimination among variants with AF = 0 (ROC-AUC = 0.9373), indicating that substantial discriminatory information remains among variants with AF = 0. Protein-sequence and structure-derived information remains an important source of signal for missense pathogenicity prediction, as illustrated by the strong performance of specialized predictors such as AlphaMissense on missense-focused benchmarks. AGVarPred does not explicitly incorporate the same protein-sequence or structure representations, although its VEP-derived SIFT and PolyPhen annotations provide protein-level consequence information. Consistent with this, gnomAD AF remained the dominant individual feature by SHAP importance, and the ClinVar development set showed strong enrichment of pathogenic variants among AF = 0 variants. The AF-stratified results therefore support a complementary role for AlphaGenome-derived functional information while not eliminating the possibility of ascertainment effects associated with population frequency and ClinVar labeling. Second, the tissue and assay SHAP rankings are descriptive. They reflect which AlphaGenome tracks were selected and how the model used them, but they should not be read as evidence of causal tissue-specific pathogenicity mechanisms. Third, the MAVE Independent result is not evidence of a flawed benchmark. It demonstrates that molecular functional impact and clinical pathogenicity are related but distinct concepts: a variant can disrupt a molecular assay without being clinically pathogenic, and clinically pathogenic variants may not produce strong assay phenotypes.

The no-allele-frequency model retains high internal performance (AUC 0.9640), indicating that AGVarPred can still discriminate pathogenic from benign variants when population frequency is unavailable. However, this model has a much higher false-positive rate on common benign variants (FPR 0.037 versus 0.0024 for the full model), confirming that allele frequency is essential for recognizing benign common variation (Supplementary Figures S2 and S5). External performance of the no-AF model was not systematically benchmarked and remains a limitation.

In sum, AGVarPred offers a useful complement to existing predictors: it scores variants across the evaluated SNV and indel sets, performs well on several clinically curated benchmarks, and provides well-calibrated probabilities on the internal ClinVar-like test distribution. However, external calibration analyses show that these probabilities do not necessarily transport to datasets with different pathogenic prevalence and label distributions. AGVarPred does not outperform the best missense predictors on missense-rich benchmarks, nor does it replace expert clinical interpretation or ACMG/AMP criteria [20].

### Limitations

Several limitations should be considered. First, AGVarPred depends on AlphaGenome-derived functional scores accessed through the AlphaGenome platform, so reproduction of feature extraction depends on continued platform access. Second, the model was trained and calibrated on ClinVar, whose disease-gene ascertainment and allele-frequency composition may influence learned associations; accordingly, calibrated probabilities should be interpreted as ClinVar-distribution scores rather than universally transportable probabilities. Third, the external benchmarks differ in label source, prevalence, variant composition, and potential overlap with comparator training data, and the VIP analysis uses ClinVar-derived labels. Fourth, AGVarPred was evaluated on the variant classes represented in the final feature matrices, primarily SNVs and short indels, and extension to structural variants remains outside the present scope. Finally, SHAP rankings are descriptive because AlphaGenome features are correlated, while subgroup analyses with small sample sizes should be considered exploratory. These limitations motivate future evaluation on broader independent datasets, external recalibration, and integration with additional sequence, protein, and clinical evidence.

### Future directions

Future work should focus on prospective evaluation in clinically relevant cohorts, calibration updating for target populations with different pathogenic prevalence, extension to additional variant classes including structural variants, comparison with a broader set of contemporary predictors, and development of per-use-case threshold selection strategies.

