## Supplementary files for "AlphaGenome informed machine learning for pathogenicity prediction of germline mutations"

**Supplementary Figures**

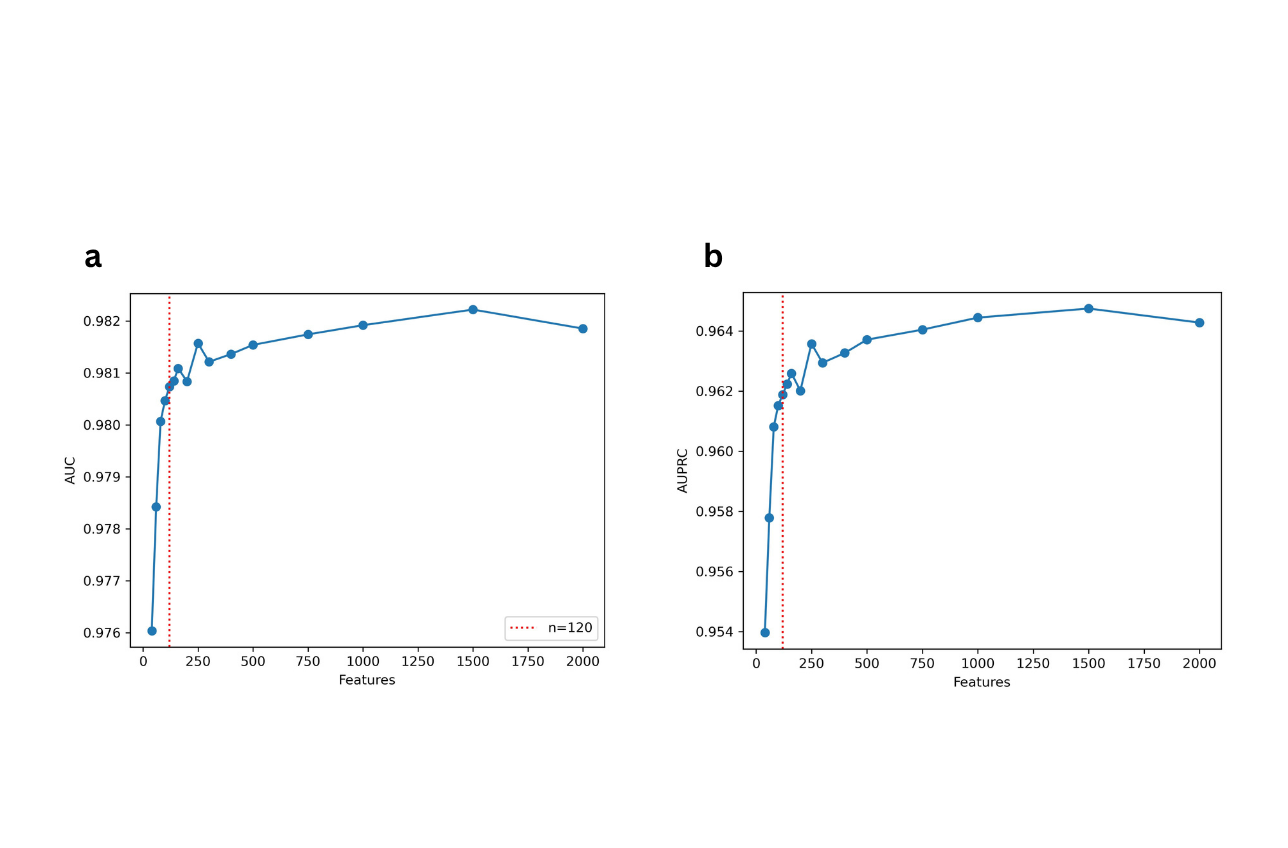

**Supplementary Figure S1.** Mean cross-validated (a) ROC-AUC and (b) PR-AUC (AUPRC) as a function of the number of top-ranked features retained during nested gene-level cross-validation. The red dotted line marks the 120-feature panel selected by the one-standard-deviation rule.

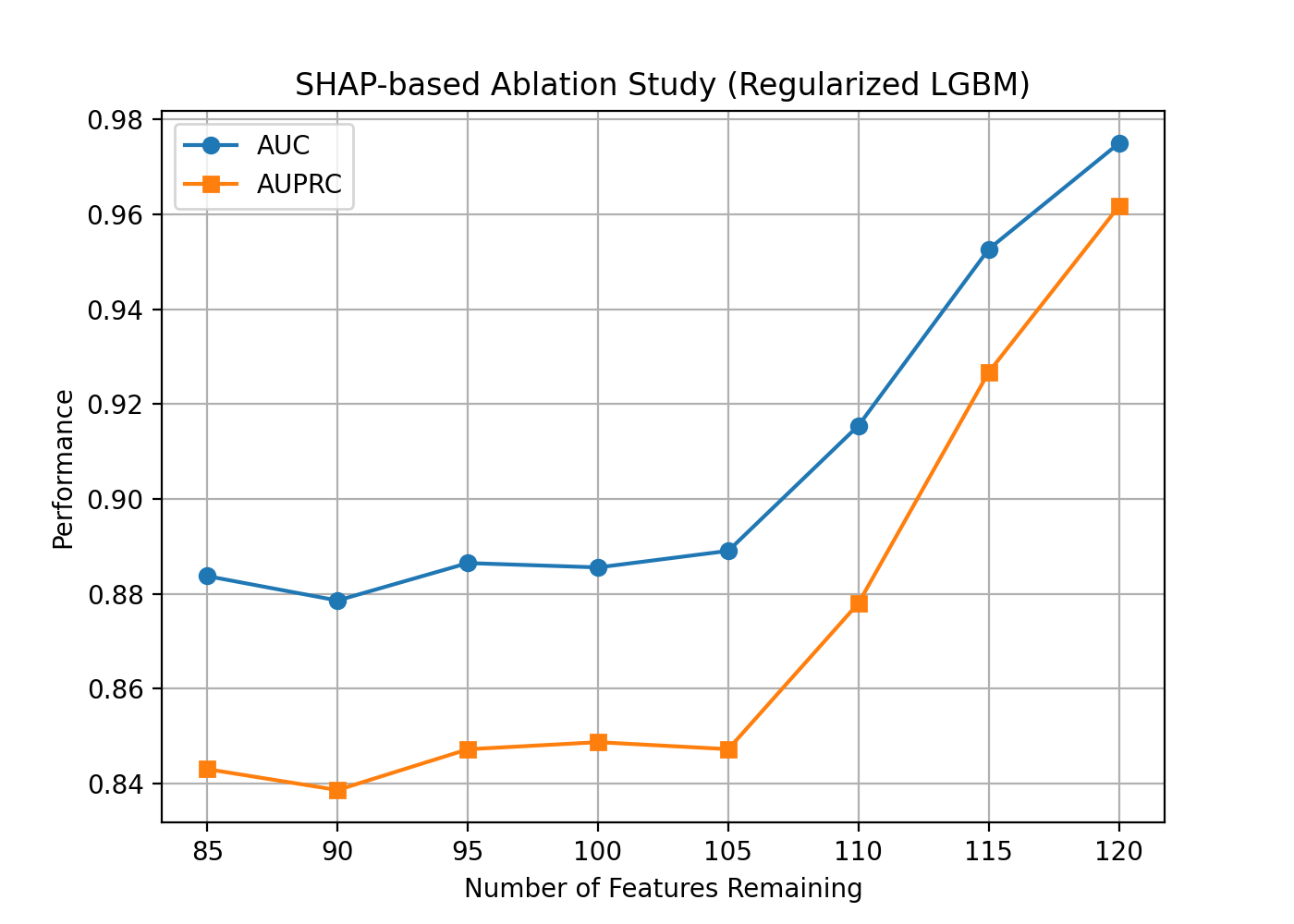

**Supplementary Figure S2.** ROC-AUC and PR-AUC (AUPRC) of the regularized LightGBM model as features are progressively removed from the full 120-feature panel.

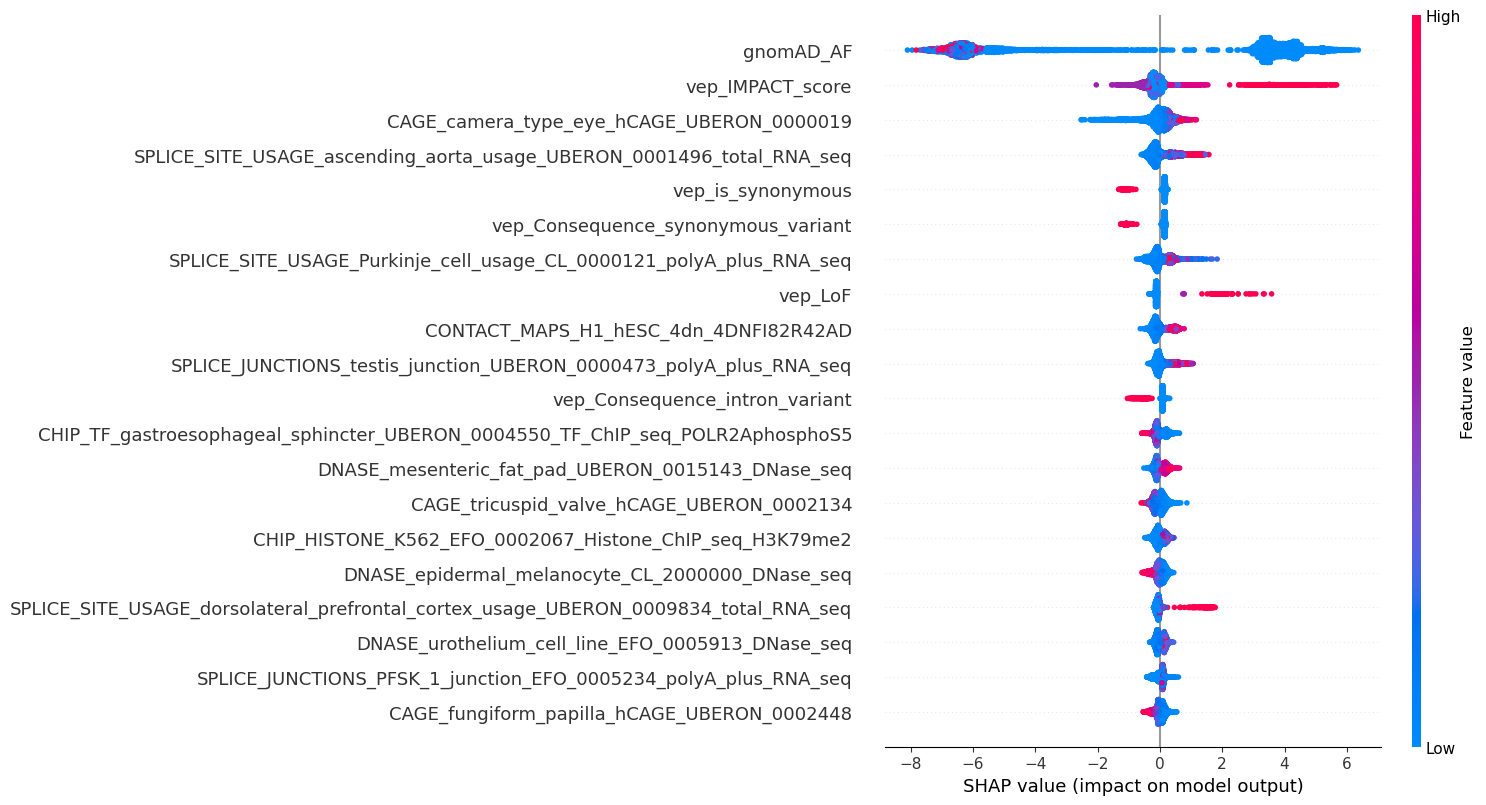

**Supplementary Figure S3.** Features of the final AGVarPred model ranked by mean absolute SHAP value; point color indicates feature value (high/low). SHAP values were computed on the uncalibrated LightGBM margin.

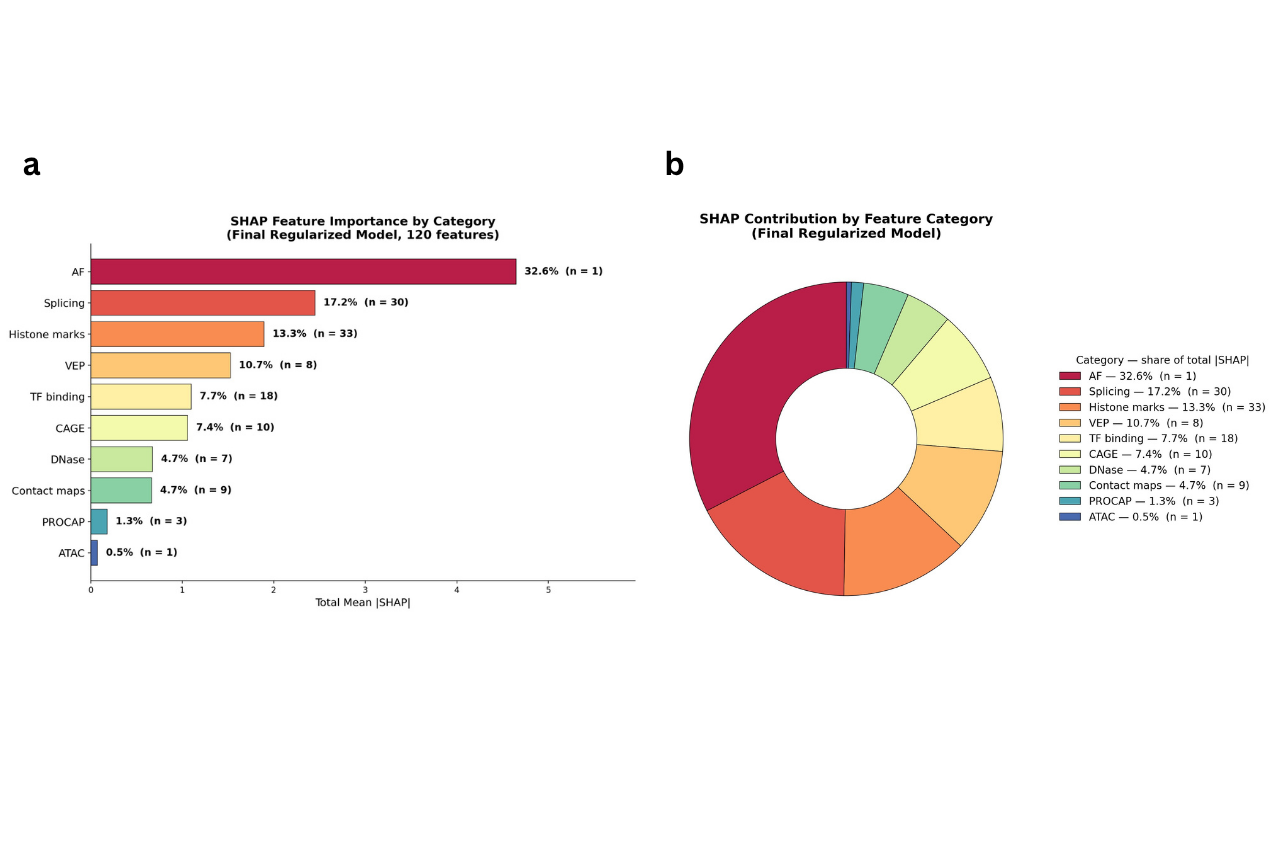

**Supplementary Figure S4.** (a) Total mean |SHAP| aggregated across feature categories for the final regularized 120-feature model; labels give each category's share of total |SHAP| and the number of contributing features (n). (b) Donut chart of the same category shares; the legend reports category, percentage of total |SHAP|, and number of features.

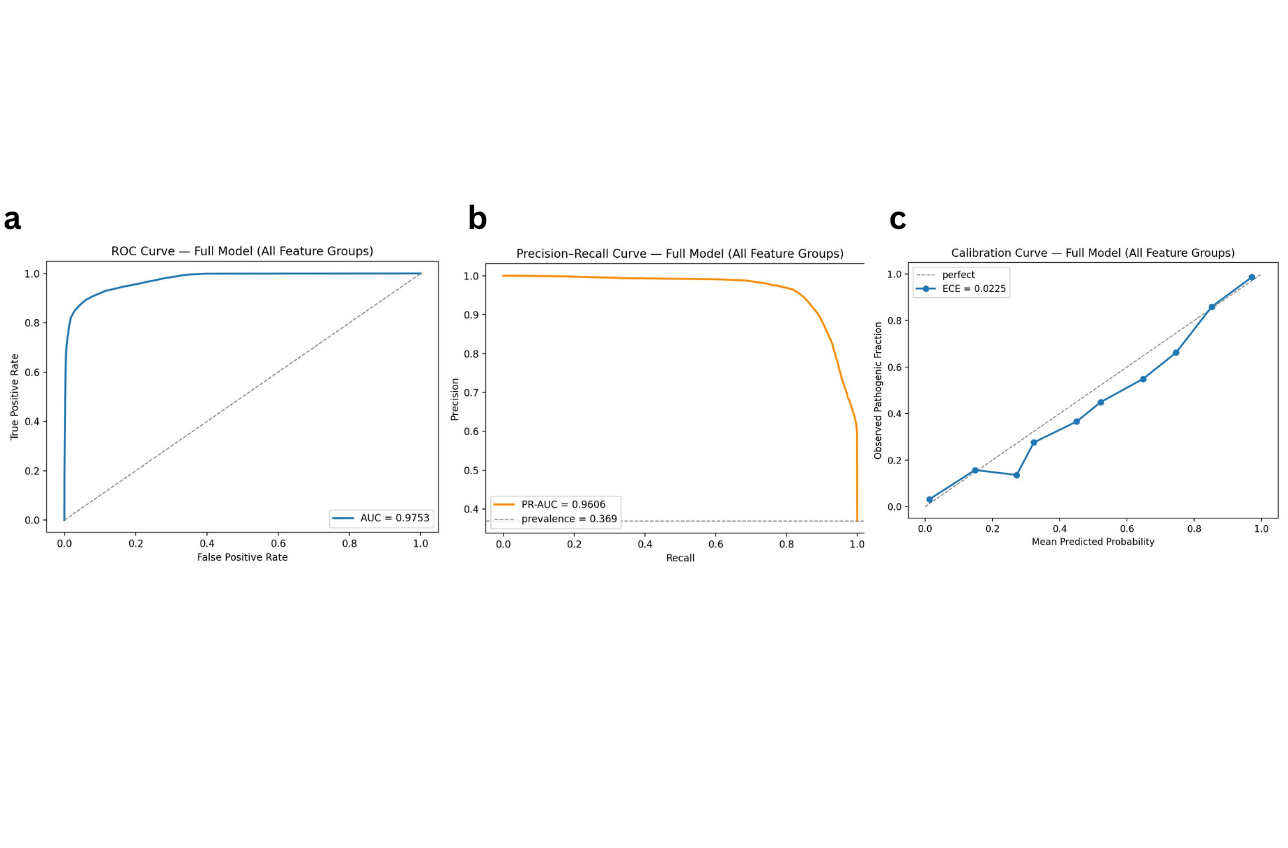

**Supplementary Figure S5.** (a) ROC, (b) precision-recall, and (c) calibration curves of the full model (all feature groups) on the 12,385 gene-held-out internal ClinVar test variants (ROC-AUC = 0.9753, PR-AUC = 0.9606, ECE = 0.0225). Curves use calibrated probabilities.

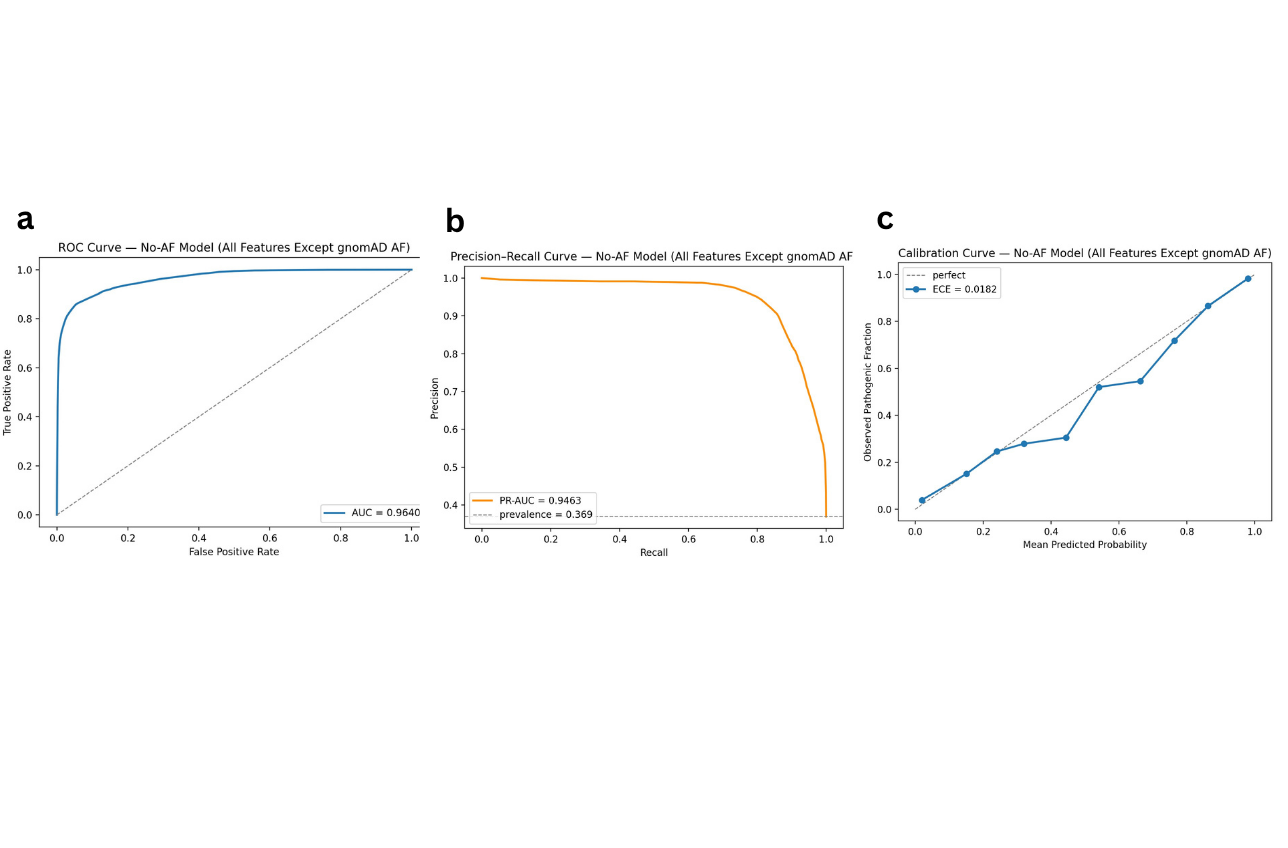

**Supplementary Figure S6.** (a) ROC, (b) precision-recall, and (c) calibration curves of the model trained without gnomAD allele-frequency features (No-AF panel) on the internal test set (ROC-AUC = 0.9640, PR-AUC = 0.9463, ECE = 0.0182). The dashed line in (b) marks the pathogenic prevalence (0.369).

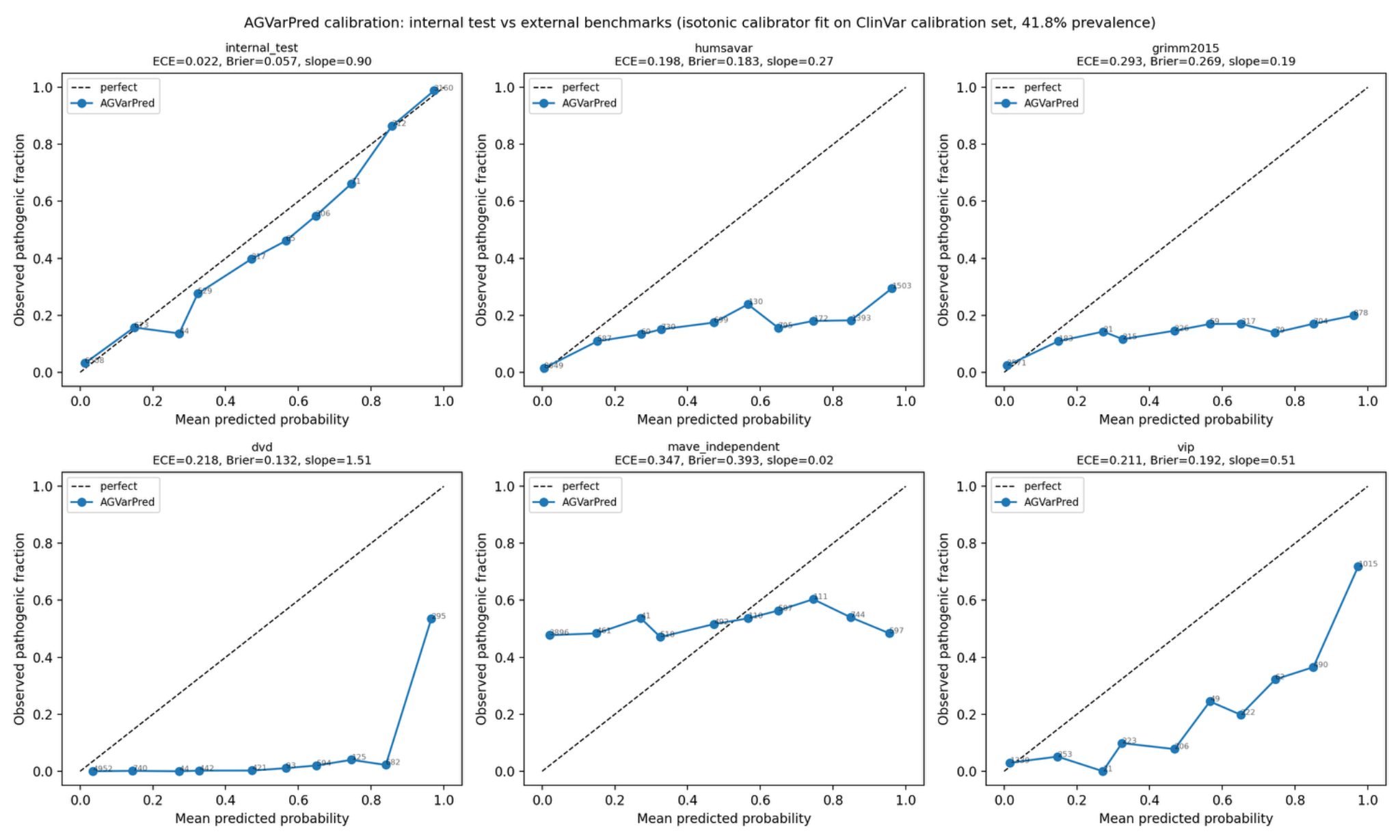

**Supplementary Figure S7.** Internal and external calibration of AGVarPred. Calibration curves for AGVarPred on the internal ClinVar test set and external benchmark datasets. The model shows good calibration on the internal ClinVar-like distribution but systematic over-confidence on several lower-prevalence external benchmarks. Calibration metrics are summarized by Brier score, 10-bin expected calibration error (ECE), calibration intercept, and calibration slope.

### **Supplementary Tables**

#### **Supplementary Table S1.** Final 120-feature panel selected by nested feature elimination

| **#** | **Feature** | **#** | **Feature** |
| --- | --- | --- | --- |
| 1 | gnomAD_AF | 61 | CHIP_HISTONE_prostate_gland_UBERON_0002367_Histone_ChIP_seq_H3K4me3 |
| 2 | vep_IMPACT_score | 62 | SPLICE_JUNCTIONS_frontal_cortex_junction_UBERON_0001870_total_RNA_seq |
| 3 | SPLICE_SITE_USAGE_muscle_of_arm_usage_UBERON_0001499_polyA_plus_RNA_seq | 63 | CHIP_HISTONE_stomach_UBERON_0000945_Histone_ChIP_seq_H3K27me3 |
| 4 | SPLICE_SITE_USAGE_ascending_aorta_usage_UBERON_0001496_total_RNA_seq | 64 | CAGE_fungiform_papilla_hCAGE_UBERON_0002448 |
| 5 | SPLICE_SITE_USAGE_Purkinje_cell_usage_CL_0000121_polyA_plus_RNA_seq | 65 | CHIP_HISTONE_muscle_of_trunk_UBERON_0001774_Histone_ChIP_seq_H3K9me3 |
| 6 | CAGE_camera_type_eye_hCAGE_UBERON_0000019 | 66 | CHIP_HISTONE_substantia_nigra_UBERON_0002038_Histone_ChIP_seq_H3K4me3 |
| 7 | SPLICE_SITE_USAGE_dorsolateral_prefrontal_cortex_usage_UBERON_0009834_total_RNA_seq | 67 | CHIP_TF_HEK293_EFO_0001182_TF_ChIP_seq_HOXB7_genetically_modified_insertion_using_CRISPR_targeting_H_sapiens_HOXB7 |
| 8 | SPLICE_SITE_USAGE_testis_usage_UBERON_0000473_total_RNA_seq | 68 | SPLICE_JUNCTIONS_HeLa_S3_junction_EFO_0002791_polyA_plus_RNA_seq |
| 9 | SPLICE_SITE_USAGE_cerebellum_usage_UBERON_0002037_total_RNA_seq | 69 | DNASE_A549_EFO_0001086_DNase_seq |
| 10 | SPLICE_SITE_USAGE_urinary_bladder_usage_UBERON_0001255_total_RNA_seq | 70 | CHIP_HISTONE_esophagus_squamous_epithelium_UBERON_0006920_Histone_ChIP_seq_H3K27me3 |
| 11 | SPLICE_JUNCTIONS_dorsolateral_prefrontal_cortex_junction_UBERON_0009834_total_RNA_seq | 71 | DNASE_COLO829_EFO_0002140_DNase_seq |
| 12 | vep_LoF | 72 | CHIP_TF_HepG2_EFO_0001187_TF_ChIP_seq_TAF1 |
| 13 | SPLICE_SITE_USAGE_metanephros_usage_UBERON_0000081_total_RNA_seq | 73 | CHIP_HISTONE_amnion_UBERON_0000305_Histone_ChIP_seq_H3K36me3 |
| 14 | CONTACT_MAPS_H1_hESC_4dn_4DNFI82R42AD | 74 | DNASE_urothelium_cell_line_EFO_0005913_DNase_seq |
| 15 | SPLICE_JUNCTIONS_BE2C_junction_EFO_0005725_polyA_plus_RNA_seq | 75 | CHIP_HISTONE_MM1S_EFO_0005724_Histone_ChIP_seq_H3K27me3 |
| 16 | vep_PolyPhen_score | 76 | CHIP_HISTONE_neuron_CL_0000540_Histone_ChIP_seq_H3K36me3 |
| 17 | SPLICE_SITES_nan_acceptor | 77 | SPLICE_JUNCTIONS_occipital_lobe_junction_UBERON_0002021_total_RNA_seq |
| 18 | SPLICE_JUNCTIONS_ascending_aorta_junction_UBERON_0001496_total_RNA_seq | 78 | CHIP_TF_A549_EFO_0001086_TF_ChIP_seq_NR3C1 |
| 19 | SPLICE_JUNCTIONS_PFSK_1_junction_EFO_0005234_polyA_plus_RNA_seq | 79 | CHIP_TF_A549_EFO_0001086_TF_ChIP_seq_REST |
| 20 | CONTACT_MAPS_skin_fibroblast_4dn_4DNFIMDOXUT8 | 80 | PROCAP_endothelial_cell_of_umbilical_vein_ENCSR801ECP |
| 21 | CHIP_HISTONE_iPS_18a_EFO_0007105_Histone_ChIP_seq_H3K9me3 | 81 | vep_Consequence_synonymous_variant |
| 22 | SPLICE_SITE_USAGE_parietal_lobe_usage_UBERON_0001872_total_RNA_seq | 82 | CHIP_HISTONE_HEK293_EFO_0001182_Histone_ChIP_seq_H3K36me3 |
| 23 | SPLICE_JUNCTIONS_testis_junction_UBERON_0000473_polyA_plus_RNA_seq | 83 | CHIP_HISTONE_VCaP_EFO_0007752_Histone_ChIP_seq_H3K27ac |
| 24 | CAGE_pancreas_hCAGE_UBERON_0001264 | 84 | CHIP_HISTONE_Panc1_EFO_0002713_Histone_ChIP_seq_H3K36me3 |
| 25 | CHIP_HISTONE_neuronal_stem_cell_CL_0000047_Histone_ChIP_seq_H3K9me3 | 85 | CAGE_neutrophil_LQhCAGE_CL_0000775 |
| 26 | SPLICE_JUNCTIONS_spinal_cord_junction_UBERON_0002240_total_RNA_seq | 86 | CHIP_HISTONE_fibroblast_of_lung_CL_0002553_Histone_ChIP_seq_H3K79me2 |
| 27 | SPLICE_JUNCTIONS_Purkinje_cell_junction_CL_0000121_polyA_plus_RNA_seq | 87 | CHIP_HISTONE_muscle_of_leg_UBERON_0001383_Histone_ChIP_seq_H3K9me3 |
| 28 | CONTACT_MAPS_HCT116_4dn_4DNFILP99QJS | 88 | ATAC_central_memory_CD8_positive_alpha_beta_T_cell_CL_0000907_ATAC_seq |
| 29 | PROCAP_MCF_10A_ENCSR860TYZ | 89 | DNASE_mesenteric_fat_pad_UBERON_0015143_DNase_seq |
| 30 | vep_SIFT_score | 90 | CAGE_umbilical_cord_blood_LQhCAGE_UBERON_0012168 |
| 31 | vep_is_synonymous | 91 | CONTACT_MAPS_H1_hESC_4dn_4DNFINFQV712 |
| 32 | CONTACT_MAPS_CyT49_4dn_4DNFI2VYSX9M | 92 | CHIP_TF_HepG2_EFO_0001187_TF_ChIP_seq_SYNCRIP |
| 33 | CAGE_parotid_gland_hCAGE_UBERON_0001831 | 93 | CONTACT_MAPS_H9_4dn_4DNFIT5YVTLO |
| 34 | CONTACT_MAPS_H9_4dn_4DNFIRO3DQCO | 94 | CHIP_TF_HeLa_S3_EFO_0002791_TF_ChIP_seq_E2F1 |
| 35 | CONTACT_MAPS_H1_hESC_4dn_4DNFI6HDY7WZ | 95 | CHIP_HISTONE_neutrophil_CL_0000775_Histone_ChIP_seq_H3K36me3 |
| 36 | SPLICE_SITES_nan_donor | 96 | CHIP_TF_HepG2_EFO_0001187_TF_ChIP_seq_ZNF384 |
| 37 | CAGE_seminal_vesicle_hCAGE_UBERON_0000998 | 97 | CHIP_TF_K562_EFO_0002067_TF_ChIP_seq_ATF1_genetically_modified_insertion_using_CRISPR_targeting_H_sapiens_ATF1 |
| 38 | DNASE_epidermal_melanocyte_CL_2000000_DNase_seq | 98 | CHIP_HISTONE_effector_memory_CD4_positive_alpha_beta_T_cell_CL_0000905_Histone_ChIP_seq_H3K27ac |
| 39 | CHIP_TF_HepG2_EFO_0001187_TF_ChIP_seq_RBFOX2 | 99 | vep_Consequence_intron_variant |
| 40 | CAGE_neuron_hCAGE_CL_0000540 | 100 | PROCAP_Calu3_ENCSR797DEF |
| 41 | CHIP_HISTONE_H9_EFO_0003045_Histone_ChIP_seq_H3K9me3 | 101 | SPLICE_JUNCTIONS_diencephalon_junction_UBERON_0001894_total_RNA_seq |
| 42 | CHIP_TF_HepG2_EFO_0001187_TF_ChIP_seq_NR2F1_genetically_modified_insertion_using_CRISPR_targeting_H_sapiens_NR2F1 | 102 | vep_Consequence_downstream_gene_variant |
| 43 | SPLICE_JUNCTIONS_kidney_junction_UBERON_0002113_polyA_plus_RNA_seq | 103 | CHIP_HISTONE_HepG2_EFO_0001187_Histone_ChIP_seq_H3K27ac |
| 44 | SPLICE_JUNCTIONS_testis_junction_UBERON_0000473_total_RNA_seq | 104 | SPLICE_JUNCTIONS_spinal_cord_junction_UBERON_0002240_polyA_plus_RNA_seq |
| 45 | CHIP_TF_HepG2_EFO_0001187_TF_ChIP_seq_ZC3H13_genetically_modified_insertion_using_CRISPR_targeting_H_sapiens_ZC3H13 | 105 | CHIP_HISTONE_neural_progenitor_cell_CL_0011020_Histone_ChIP_seq_H4K20me1 |
| 46 | CHIP_TF_HepG2_EFO_0001187_TF_ChIP_seq_POLR2AphosphoS5 | 106 | SPLICE_SITE_USAGE_testis_usage_UBERON_0000473_gtex_Testis_polyA_plus_RNA_seq |
| 47 | CAGE_vertebral_mesenchymal_stem_cell_hCAGE_CL_0002572 | 107 | DNASE_SW480_EFO_0002083_DNase_seq |
| 48 | SPLICE_JUNCTIONS_motor_neuron_junction_CL_0000100_total_RNA_seq | 108 | CHIP_HISTONE_peripheral_blood_mononuclear_cell_CL_2000001_Histone_ChIP_seq_H3K4me3 |
| 49 | DNASE_chondrocyte_CL_0000138_DNase_seq | 109 | CHIP_TF_gastroesophageal_sphincter_UBERON_0004550_TF_ChIP_seq_POLR2AphosphoS5 |
| 50 | CHIP_HISTONE_hepatocyte_CL_0000182_Histone_ChIP_seq_H3K79me2 | 110 | CHIP_TF_SK_N_SH_EFO_0003072_TF_ChIP_seq_FOXJ3_genetically_modified_insertion_using_CRISPR_targeting_H_sapiens_FOXJ3 |
| 51 | CHIP_TF_HepG2_EFO_0001187_TF_ChIP_seq_NR2C2 | 111 | SPLICE_SITE_USAGE_cerebellum_usage_UBERON_0002037_polyA_plus_RNA_seq |
| 52 | CHIP_HISTONE_K562_EFO_0002067_Histone_ChIP_seq_H3K79me2 | 112 | CHIP_HISTONE_HepG2_EFO_0001187_Histone_ChIP_seq_H4K20me1 |
| 53 | SPLICE_SITE_USAGE_BE2C_usage_EFO_0005725_polyA_plus_RNA_seq | 113 | CHIP_HISTONE_skin_epidermis_UBERON_0001003_Histone_ChIP_seq_H3K27ac |
| 54 | CONTACT_MAPS_CyT49_4dn_4DNFIK3NF6R2 | 114 | CHIP_HISTONE_SK_N_SH_EFO_0003072_Histone_ChIP_seq_H4K20me1 |
| 55 | CHIP_HISTONE_neural_cell_CL_0002319_Histone_ChIP_seq_H3K79me2 | 115 | CHIP_HISTONE_PC_9_EFO_0002847_Histone_ChIP_seq_H3F3A |
| 56 | CHIP_HISTONE_prostate_gland_UBERON_0002367_Histone_ChIP_seq_H3K36me3 | 116 | CHIP_TF_SK_N_SH_EFO_0003072_TF_ChIP_seq_ZBTB33 |
| 57 | CHIP_TF_K562_EFO_0002067_TF_ChIP_seq_BRF2 | 117 | CAGE_tricuspid_valve_hCAGE_UBERON_0002134 |
| 58 | CHIP_HISTONE_brain_UBERON_0000955_Histone_ChIP_seq_H3K27me3 | 118 | CHIP_HISTONE_PC_9_EFO_0002847_Histone_ChIP_seq_H3K9me2 |
| 59 | CHIP_HISTONE_HUES48_EFO_0007083_Histone_ChIP_seq_H3K9me3 | 119 | CHIP_HISTONE_cardiac_muscle_cell_CL_0000746_Histone_ChIP_seq_H3K36me3 |
| 60 | CHIP_TF_HEK293_EFO_0001182_TF_ChIP_seq_ZSCAN4_genetically_modified_insertion_using_site_specific_recombination_targeting_H_sapiens_ZSCAN4 | 120 | SPLICE_SITE_USAGE_type_B_pancreatic_cell_usage_CL_0000169_total_RNA_seq |

Note. Features are listed in selection order from feature_selection_output_nested/selected_features.txt. The panel comprises 111 AlphaGenome functional-score features, one gnomAD allele-frequency feature (gnomAD_AF), and eight VEP-derived features.

#### **Supplementary Table S2.** Optuna search space and final LightGBM hyperparameters

| **Parameter** | **Search space** | **Final value** |
| --- | --- | --- |
| n_estimators | 200-500 (int) | 485 |
| learning_rate | 0.01-0.08 | 0.0653 |
| num_leaves | 20-80 (int) | 73 |
| max_depth | 3-8 (int) | 8 |
| min_child_samples | 10-100 (int) | 91 |
| subsample | 0.6-1.0 | 0.6013 |
| colsample_bytree | 0.6-1.0 | 0.6757 |
| reg_alpha | 1e-3-10.0 (log) | 0.0373 |
| reg_lambda | 1e-3-10.0 (log) | 0.0140 |
| scale_pos_weight | fixed | 2.2116 |
| random_state | fixed | 42 |
| n_jobs | fixed | -1 |

Note. Optuna TPE sampler, seed 42, 100 trials; objective was mean five-fold gene-group AUPRC. Best trial: #64 (CV AUPRC 0.9676). The no-allele-frequency model was trained analogously on 119 features.

#### **Supplementary Table S3.** Pairwise DeLong test results with FDR correction

| **Benchmark** | **Comparison** | **N common** | **N path.** | **N benign** | **AUC AGVarPred** | **AUC competitor** | **ΔAUC (95% CI)** | **q value** | **Significant** |
| --- | --- | --- | --- | --- | --- | --- | --- | --- | --- |
| humsavar | CADD | 14,716 | 1316 | 13400 | 0.826 | 0.871 | -0.045 (-0.056 to -0.034) | 2.3e-14 | yes |
| humsavar | AlphaMissense | 4,707 | 319 | 4388 | 0.833 | 0.882 | -0.049 (-0.073 to -0.025) | 0.0001 | yes |
| humsavar | REVEL | 14,379 | 1297 | 13082 | 0.830 | 0.942 | -0.112 (-0.122 to -0.102) | q < 0.001 | yes |
| mave_independent | CADD | 6,663 | 3326 | 3337 | 0.529 | 0.562 | -0.033 (-0.048 to -0.018) | 2.9e-05 | yes |
| mave_independent | AlphaMissense | 483 | 250 | 233 | 0.535 | 0.697 | -0.161 (-0.233 to -0.090) | 2.4e-05 | yes |
| mave_independent | REVEL | 2,299 | 1188 | 1111 | 0.501 | 0.699 | -0.199 (-0.229 to -0.168) | q < 0.001 | yes |
| gnomad_benign | CADD | 4,989 | 0 | 4989 | — | — | n/a | n/a | no (single-class subset) |
| gnomad_benign | AlphaMissense | 516 | 0 | 516 | — | — | n/a | n/a | no (single-class subset) |
| gnomad_benign | REVEL | 1,235 | 0 | 1235 | — | — | n/a | n/a | no (single-class subset) |
| grimm2015 | CADD | 3,815 | 30 | 3785 | 0.717 | 0.695 | 0.022 (-0.045 to 0.089) | 0.531 | no |
| grimm2015 | AlphaMissense | 607 | 0 | 607 | — | — | n/a | n/a | no (single-class subset) |
| grimm2015 | REVEL | 5,183 | 509 | 4679 | 0.746 | 0.641 | 0.105 (0.081 to 0.130) | q < 0.001 | yes |
| dvd | CADD | 8,279 | 135 | 8144 | 0.975 | 0.997 | -0.022 (-0.034 to -0.011) | 0.0003 | yes |
| dvd | AlphaMissense | 143 | 4 | 139 | 0.716 | 0.978 | -0.263 (-0.624 to 0.098) | 0.1693 | no |
| dvd | REVEL | 855 | 63 | 792 | 0.903 | 0.979 | -0.076 (-0.120 to -0.032) | 0.0009 | yes |

Note. Comparisons marked n/a were not applicable because the common subset contained variants of only one class. q values are Benjamini-Hochberg FDR-adjusted across the 11 valid pairwise tests; significance at q < 0.05.

**Supplementary Table S4.** Internal ClinVar test-set performance by variant consequence class

| **Consequence class** | **N** | **N path.** | **N benign** | **ROC-AUC (95% CI)** | **PR-AUC** | **MCC** |
| --- | --- | --- | --- | --- | --- | --- |
| missense | 957 | 147 | 810 | 0.9842 (0.9703-0.9947) | 0.9472 | 0.8457 |
| synonymous | 1,230 | 0 | 1230 | n/a | n/a | n/a |
| stop gained | 341 | 338 | 3 | 1.0000 (1.0000-1.0000) | 1.0000 | 1.0000 |
| frameshift | 293 | 291 | 2 | 0.9983 (0.9914-1.0000) | 1.0000 | 0.4966 |
| splice | 361 | 86 | 275 | 0.9997 (0.9989-1.0000) | 0.9989 | 0.9704 |
| UTR | 176 | 27 | 149 | 1.0000 (1.0000-1.0000) | 1.0000 | 1.0000 |
| intron | 1,622 | 9 | 1613 | 0.9995 (0.9985-1.0000) | 0.8963 | 0.8652 |
| non coding exon | 53 | 10 | 43 | 1.0000 (1.0000-1.0000) | 1.0000 | 1.0000 |
| other | 7,352 | 3668 | 3684 | 0.9413 (0.9358-0.9466) | 0.9453 | 0.7591 |

Note. Classes marked n/a contained no pathogenic variants (or no benign variants), so ROC-AUC could not be computed. Estimates from classes with very small negative sets (e.g., stop-gained, frameshift) are illustrative only.

#### **Supplementary Table S5.** External benchmark performance of AGVarPred by variant consequence class

| **Benchmark** | **Consequence class** | **N** | **N path.** | **ROC-AUC (95% CI)** | **PR-AUC** | **MCC** |
| --- | --- | --- | --- | --- | --- | --- |
| humsavar | missense | 8,817 | 461 | 0.911 (0.899-0.923) | 0.373 | 0.364 |
| humsavar | other | 4,968 | 813 | 0.644 (0.626-0.662) | 0.225 | 0.151 |
| humsavar | non coding exon | 137 | 4 | 0.730 (0.440-0.925) | 0.102 | 0.113 |
| humsavar | UTR | 182 | 8 | 0.842 (0.677-0.969) | 0.335 | 0.288 |
| humsavar | intron | 359 | 11 | 0.900 (0.845-0.946) | 0.186 | 0.252 |
| humsavar | splice | 198 | 16 | 0.939 (0.902-0.972) | 0.495 | 0.449 |
| humsavar | synonymous | 25 | 3 | 0.591 (0.250-0.909) | 0.197 | 0.123 |
| humsavar | stop gained | 32 | 1 | 0.839 (0.742-0.919) | 0.100 | 0.046 |
| mave_independent | other | 7,388 | 3690 | 0.532 (0.520-0.545) | 0.518 | 0.054 |
| mave_independent | missense | 104 | 35 | 0.568 (0.448-0.676) | 0.389 | 0.155 |
| mave_independent | synonymous | 65 | 26 | 0.427 (0.280-0.567) | 0.359 | -0.199 |
| gnomad_benign | UTR | 322 | 0 | n/a | n/a | n/a |
| gnomad_benign | synonymous | 794 | 0 | n/a | n/a | n/a |
| gnomad_benign | intron | 1,812 | 0 | n/a | n/a | n/a |
| gnomad_benign | missense | 784 | 0 | n/a | n/a | n/a |
| gnomad_benign | splice | 173 | 0 | n/a | n/a | n/a |
| gnomad_benign | stop gained | 25 | 0 | n/a | n/a | n/a |
| gnomad_benign | other | 829 | 0 | n/a | n/a | n/a |
| gnomad_benign | non coding exon | 250 | 0 | n/a | n/a | n/a |
| grimm2015 | missense | 4,382 | 409 | 0.746 (0.727-0.766) | 0.185 | 0.226 |
| grimm2015 | splice | 81 | 10 | 0.722 (0.528-0.872) | 0.265 | 0.183 |
| grimm2015 | other | 577 | 87 | 0.718 (0.673-0.760) | 0.239 | 0.234 |
| grimm2015 | intron | 127 | 4 | 0.741 (0.468-0.949) | 0.113 | 0.186 |
| grimm2015 | synonymous | 39 | 1 | 0.145 (0.053-0.250) | 0.029 | -0.055 |
| grimm2015 | non coding exon | 24 | 0 | n/a | n/a | n/a |
| grimm2015 | UTR | 23 | 1 | 0.727 (0.545-0.909) | 0.143 | -0.127 |
| dvd | UTR | 307 | 0 | n/a | n/a | n/a |
| dvd | missense | 725 | 59 | 0.939 (0.906-0.967) | 0.701 | 0.295 |
| dvd | synonymous | 267 | 0 | n/a | n/a | n/a |
| dvd | splice | 67 | 22 | 0.930 (0.829-1.000) | 0.940 | 0.244 |
| dvd | intron | 6,792 | 1 | 1.000 (1.000-1.000) | 1.000 | 0.026 |
| dvd | stop gained | 39 | 39 | n/a | n/a | n/a |
| dvd | frameshift | 54 | 54 | n/a | n/a | n/a |
| dvd | other | 137 | 20 | 0.513 (0.368-0.669) | 0.297 | 0.093 |

Note. Estimates from subgroups with fewer than 100 pathogenic variants, fewer than 100 total variants, or fewer than 10 pathogenic variants are exploratory. gnomAD Common Benign subgroups contain only benign variants, so ROC-AUC is not applicable.

#### **Supplementary Table S6.** Non-missense ROC-AUC comparison on the AGVarPred-scored universe

| **Benchmark** | **Tool** | **N scored** | **N path.** | **ROC-AUC (95% CI)** | **PR-AUC (95% CI)** |
| --- | --- | --- | --- | --- | --- |
| humsavar | AGVarPred | 5,901 | 856 | 0.683 (0.668-0.698) | 0.220 (0.207-0.235) |
| humsavar | CADD | 5,899 | 855 | 0.849 (0.836-0.862) | 0.412 (0.389-0.438) |
| humsavar | AlphaMissense | 1,867 | 211 | 0.882 (0.854-0.908) | 0.580 (0.517-0.649) |
| humsavar | REVEL | 5,583 | 836 | 0.935 (0.925-0.942) | 0.723 (0.694-0.753) |
| mave_independent | AGVarPred | 7,453 | 3716 | 0.532 (0.519-0.545) | 0.516 (0.506-0.528) |
| mave_independent | CADD | 6,559 | 3291 | 0.564 (0.551-0.578) | 0.563 (0.551-0.577) |
| mave_independent | AlphaMissense | 466 | 246 | 0.688 (0.635-0.737) | 0.666 (0.618-0.723) |
| mave_independent | REVEL | 2,195 | 1153 | 0.698 (0.677-0.721) | 0.705 (0.680-0.732) |
| gnomad_benign | AGVarPred | 4,205 | 0 | n/a | n/a |
| gnomad_benign | CADD | 4,205 | 0 | n/a | n/a |
| gnomad_benign | AlphaMissense | 209 | 0 | n/a | n/a |
| gnomad_benign | REVEL | 463 | 0 | n/a | n/a |
| grimm2015 | AGVarPred | 871 | 103 | 0.7516 (0.7101-0.7884) | 0.222 (0.189-0.268) |
| grimm2015 | CADD | 591 | 2 | 0.7322  (0.633-0.821) | 0.009 (0.008-0.018) |
| grimm2015 | AlphaMissense | 93 | 0 | n/a | n/a |
| grimm2015 | REVEL | 821 | 100 | 0.700 (0.638-0.751) | 0.268 (0.210-0.345) |
| dvd | AGVarPred | 7,663 | 136 | 0.981 (0.970-0.990) | 0.836 (0.783-0.886) |
| dvd | CADD | 7,555 | 77 | 0.998 (0.996-1.000) | 0.952 (0.910-0.987) |
| dvd | AlphaMissense | 73 | 2 | 0.979 (0.930-1.000) | 0.700 (0.286-1.000) |
| dvd | REVEL | 131 | 5 | 0.992 (0.975-1.000) | 0.858 (0.626-1.000) |

Note. Evaluations are restricted to non-missense variants within the AGVarPred-scored universe; each competitor is reported on the subset it could score. gnomAD Common Benign contains no pathogenic non-missense variants.

#### **Supplementary Table S7.** SHAP importance aggregated by tissue group

| **Tissue group** | **N features** | **N biosamples** | **Total SHAP** | **% of AlphaGenome SHAP mass** | **95% CI** |
| --- | --- | --- | --- | --- | --- |
| Brain / CNS | 26 | 19 | 1.780 | 22.0% | 1.397-2.263 |
| Liver / digestive | 23 | 13 | 1.593 | 19.7% | 1.312-1.871 |
| Heart / vasculature | 6 | 5 | 0.792 | 9.8% | 0.511-1.154 |
| Blood / immune | 10 | 8 | 0.677 | 8.4% | 0.537-0.842 |
| Reproductive | 9 | 6 | 0.656 | 8.1% | 0.452-0.909 |
| Embryonic / stem | 11 | 7 | 0.640 | 7.9% | 0.394-0.996 |
| Lung | 7 | 4 | 0.366 | 4.5% | 0.262-0.473 |
| Kidney / urinary | 4 | 4 | 0.334 | 4.1% | 0.266-0.420 |
| Other | 5 | 2 | 0.301 | 3.7% | 0.233-0.381 |
| Eye / sensory | 1 | 1 | 0.290 | 3.6% | n/a |
| Skin | 4 | 4 | 0.286 | 3.5% | 0.184-0.415 |
| Muscle | 3 | 3 | 0.201 | 2.5% | 0.090-0.319 |
| Connective / mesenchyme | 2 | 2 | 0.180 | 2.2% | 0.168-0.192 |

Note. Percentages are relative to the total SHAP mass of the 111 AlphaGenome features. Bootstrap confidence intervals were computed by resampling features within each group; single-feature groups have no interval (n/a).

#### **Supplementary Table S8.** Decision-curve analysis summary

| **Benchmark** | **Tool** | **N scored** | **Prevalence** | **Max net benefit (threshold)** | **Integrated net benefit** | **Positive-NB range** | **Beats-both range** |
| --- | --- | --- | --- | --- | --- | --- | --- |
| dvd | AGVarPred | 8,388 | 0.023 | 0.015 (0.01) | -0.146 | 0.01-0.03 | 0.01-0.03 |
| dvd | AlphaMissense | 143 | 0.028 | 0.018 (0.01) | -0.010 | 0.01-0.99 | 0.28-0.99 |
| dvd | REVEL | 855 | 0.074 | 0.065 (0.01) | 0.033 | 0.01-0.97 | 0.01-0.97 |
| gnomad_benign | AGVarPred | 4,989 | 0.000 | 0.000 (0.8300000000000001) | -0.001 | none | none |
| gnomad_benign | AlphaMissense | 516 | 0.000 | -0.010 (0.01) | -0.063 | none | none |
| gnomad_benign | REVEL | 1,235 | 0.000 | 0.000 (0.98) | -0.041 | none | none |
| grimm2015 | AGVarPred | 5,253 | 0.097 | 0.086 (0.01) | -0.437 | 0.01-0.17 | 0.02-0.17 |
| grimm2015 | AlphaMissense | 607 | 0.000 | -0.010 (0.01) | -0.121 | none | none |
| grimm2015 | REVEL | 5,183 | 0.098 | 0.089 (0.01) | -0.045 | 0.01-0.13 | 0.06-0.13 |
| humsavar | AGVarPred | 14,718 | 0.089 | 0.082 (0.01) | -0.239 | 0.01-0.20 | 0.01-0.20 |
| humsavar | AlphaMissense | 4,707 | 0.068 | 0.058 (0.01) | -0.058 | 0.01-0.19 | 0.02-0.19 |
| humsavar | REVEL | 14,379 | 0.090 | 0.081 (0.01) | 0.015 | 0.01-0.99 | 0.01-0.99 |
| internal_test | AGVarPred | 12,385 | 0.369 | 0.366 (0.01) | 0.280 | 0.01-0.99 | 0.01-0.99 |
| mave_independent | AGVarPred | 7,557 | 0.496 | 0.362 (0.01) | -0.016 | 0.01-0.53 | 0.49-0.53 |
| mave_independent | AlphaMissense | 483 | 0.518 | 0.513 (0.01) | -0.216 | 0.01-0.60 | 0.04-0.60 |
| mave_independent | REVEL | 2,299 | 0.517 | 0.512 (0.01) | 0.169 | 0.01-0.76 | 0.01-0.76 |

Note. Positive-NB range = thresholds with net benefit > 0; beats-both range = thresholds where the tool exceeds both treat-all and treat-none strategies. CADD was excluded because phred scores are not calibrated probabilities.

#### **Supplementary Table S9.** SHAP importance by biosample

| **Biosample** | **Tissue group** | **N features** | **Total SHAP** | **% of AlphaGenome SHAP mass** | **95% CI** |
| --- | --- | --- | --- | --- | --- |
| HepG2 EFO 0001187 | Liver / digestive | 10 | 0.686 | 8.47% | 0.555-0.829 |
| ascending aorta | Heart / vasculature | 2 | 0.354 | 4.37% | 0.151-0.556 |
| testis | Reproductive | 4 | 0.320 | 3.95% | 0.140-0.558 |
| H1 hESC | Embryonic / stem | 3 | 0.298 | 3.69% | 0.070-0.604 |
| camera type eye | Eye / sensory | 1 | 0.290 | 3.58% | n/a |
| Purkinje cell | Brain / CNS | 2 | 0.262 | 3.23% | 0.074-0.450 |
| K562 EFO 0002067 | Blood / immune | 3 | 0.233 | 2.88% | 0.101-0.361 |
| dorsolateral prefrontal cortex | Brain / CNS | 2 | 0.204 | 2.52% | 0.174-0.234 |
| BE2C | Brain / CNS | 2 | 0.190 | 2.35% | 0.182-0.198 |
| SK N SH EFO 0003072 | Brain / CNS | 3 | 0.159 | 1.96% | 0.072-0.274 |
| Unassigned | Other | 2 | 0.158 | 1.96% | 0.143-0.174 |
| A549 EFO 0001086 | Lung | 3 | 0.153 | 1.89% | 0.091-0.226 |
| gastroesophageal sphincter UBERON 0004550 | Liver / digestive | 1 | 0.144 | 1.78% | n/a |
| mesenteric fat pad UBERON 0015143 | Heart / vasculature | 1 | 0.143 | 1.76% | n/a |
| HEK293 EFO 0001182 | Other | 3 | 0.142 | 1.76% | 0.127-0.155 |
| tricuspid valve | Heart / vasculature | 1 | 0.141 | 1.75% | n/a |
| H9 | Embryonic / stem | 2 | 0.136 | 1.67% | 0.105-0.166 |
| epidermal melanocyte CL 2000000 | Skin | 1 | 0.120 | 1.48% | n/a |
| PC 9 EFO 0002847 | Lung | 2 | 0.117 | 1.45% | 0.073-0.161 |
| urothelium cell line EFO 0005913 | Kidney / urinary | 1 | 0.114 | 1.41% | n/a |
| PFSK 1 | Brain / CNS | 1 | 0.109 | 1.35% | n/a |
| fungiform papilla | Liver / digestive | 1 | 0.109 | 1.35% | n/a |
| muscle of arm | Muscle | 1 | 0.106 | 1.31% | n/a |
| prostate gland UBERON 0002367 | Liver / digestive | 2 | 0.099 | 1.23% | 0.038-0.160 |
| chondrocyte CL 0000138 | Connective / mesenchyme | 1 | 0.096 | 1.19% | n/a |
| parietal lobe | Brain / CNS | 1 | 0.095 | 1.18% | n/a |
| parotid gland | Liver / digestive | 1 | 0.093 | 1.15% | n/a |
| neutrophil CL 0000775 | Blood / immune | 1 | 0.091 | 1.12% | n/a |
| cardiac muscle cell CL 0000746 | Heart / vasculature | 1 | 0.089 | 1.10% | n/a |
| MCF 10A | Reproductive | 1 | 0.086 | 1.06% | n/a |
| pancreas | Liver / digestive | 1 | 0.085 | 1.05% | n/a |
| vertebral mesenchymal stem cell | Connective / mesenchyme | 1 | 0.084 | 1.04% | n/a |
| HCT116 | Liver / digestive | 1 | 0.082 | 1.02% | n/a |
| neuron CL 0000540 | Brain / CNS | 1 | 0.081 | 1.00% | n/a |
| urinary bladder | Kidney / urinary | 1 | 0.079 | 0.98% | n/a |
| metanephros | Kidney / urinary | 1 | 0.078 | 0.97% | n/a |
| CyT49 | Embryonic / stem | 2 | 0.078 | 0.96% | 0.068-0.088 |
| substantia nigra UBERON 0002038 | Brain / CNS | 1 | 0.077 | 0.95% | n/a |
| HeLa S3 | Reproductive | 1 | 0.077 | 0.95% | n/a |
| neuron | Brain / CNS | 1 | 0.075 | 0.93% | n/a |
| brain UBERON 0000955 | Brain / CNS | 1 | 0.075 | 0.93% | n/a |
| central memory CD8 positive alpha beta T cell CL 0000907 | Blood / immune | 1 | 0.074 | 0.92% | n/a |
| skin fibroblast | Skin | 1 | 0.073 | 0.90% | n/a |
| spinal cord | Brain / CNS | 2 | 0.072 | 0.89% | 0.063-0.081 |
| motor neuron | Brain / CNS | 1 | 0.072 | 0.88% | n/a |
| SW480 EFO 0002083 | Liver / digestive | 1 | 0.071 | 0.88% | n/a |
| seminal vesicle | Reproductive | 1 | 0.070 | 0.87% | n/a |
| fibroblast of lung CL 0002553 | Lung | 1 | 0.067 | 0.83% | n/a |
| endothelial cell of umbilical vein | Heart / vasculature | 1 | 0.065 | 0.80% | n/a |
| muscle of trunk UBERON 0001774 | Muscle | 1 | 0.065 | 0.80% | n/a |
| MM1S EFO 0005724 | Blood / immune | 1 | 0.063 | 0.78% | n/a |
| kidney | Kidney / urinary | 1 | 0.062 | 0.77% | n/a |
| HeLa S3 EFO 0002791 | Reproductive | 1 | 0.060 | 0.74% | n/a |
| neutrophil | Blood / immune | 1 | 0.059 | 0.72% | n/a |
| COLO829 EFO 0002140 | Skin | 1 | 0.057 | 0.70% | n/a |
| neural progenitor cell CL 0011020 | Brain / CNS | 1 | 0.056 | 0.69% | n/a |
| peripheral blood mononuclear cell CL 2000001 | Blood / immune | 1 | 0.054 | 0.67% | n/a |
| hepatocyte CL 0000182 | Liver / digestive | 1 | 0.053 | 0.65% | n/a |
| effector memory CD4 positive alpha beta T cell CL 0000905 | Blood / immune | 1 | 0.052 | 0.64% | n/a |
| cerebellum | Brain / CNS | 2 | 0.051 | 0.63% | 0.048-0.054 |
| umbilical cord blood | Blood / immune | 1 | 0.051 | 0.63% | n/a |
| frontal cortex | Brain / CNS | 1 | 0.051 | 0.63% | n/a |
| Panc1 EFO 0002713 | Liver / digestive | 1 | 0.049 | 0.60% | n/a |
| stomach UBERON 0000945 | Liver / digestive | 1 | 0.045 | 0.56% | n/a |
| VCaP EFO 0007752 | Reproductive | 1 | 0.044 | 0.54% | n/a |
| neuronal stem cell CL 0000047 | Brain / CNS | 1 | 0.042 | 0.52% | n/a |
| amnion UBERON 0000305 | Embryonic / stem | 1 | 0.039 | 0.48% | n/a |
| type B pancreatic cell | Liver / digestive | 1 | 0.039 | 0.48% | n/a |
| neural cell CL 0002319 | Brain / CNS | 1 | 0.038 | 0.47% | n/a |
| diencephalon | Brain / CNS | 1 | 0.038 | 0.46% | n/a |
| skin epidermis UBERON 0001003 | Skin | 1 | 0.037 | 0.46% | n/a |
| esophagus squamous epithelium UBERON 0006920 | Liver / digestive | 1 | 0.037 | 0.46% | n/a |
| occipital lobe | Brain / CNS | 1 | 0.034 | 0.42% | n/a |
| iPS 18a EFO 0007105 | Embryonic / stem | 1 | 0.032 | 0.40% | n/a |
| muscle of leg UBERON 0001383 | Muscle | 1 | 0.030 | 0.37% | n/a |
| HUES48 EFO 0007083 | Embryonic / stem | 1 | 0.028 | 0.35% | n/a |
| Calu3 | Lung | 1 | 0.028 | 0.35% | n/a |
| H9 EFO 0003045 | Embryonic / stem | 1 | 0.028 | 0.35% | n/a |

Note. Biosamples are ranked by total SHAP contribution. Confidence intervals are available only for biosamples represented by more than one feature. “Unassigned” represents two SPLICE_SITES features for which the source track name did not contain a biosample identifier.

**Supplementary Table S10.** Ablation analysis of AGVarPred feature groups

| **Model** | **Features** | **Threshold** | **ROC-AUC (95% CI)** | **PR-AUC** | **Brier** | **ECE** | **Sensitivity** | **Specificity** | **F1** | **MCC** |
| --- | --- | --- | --- | --- | --- | --- | --- | --- | --- | --- |
| AF only | 1 | 0.38 | 0.8115 (0.8062-0.8171) | 0.6415 | 0.1537 | 0.0417 | 0.9884 | 0.5523 | 0.7182 | 0.5463 |
| VEP only | 8 | 0.32 | 0.8044 (0.7988-0.8102) | 0.6320 | 0.1585 | 0.0405 | 0.9670 | 0.5542 | 0.7090 | 0.5236 |
| AF + VEP | 9 | 0.24 | 0.8214 (0.8164-0.8268) | 0.6559 | 0.1479 | 0.0393 | 0.9972 | 0.5576 | 0.7246 | 0.5604 |
| AlphaGenome only | 111 | 0.33 | 0.8664 (0.8601-0.8717) | 0.8290 | 0.1277 | 0.0226 | 0.7225 | 0.8713 | 0.7440 | 0.6016 |
| AlphaGenome + VEP (no AF) | 119 | 0.45 | 0.9640 (0.9612-0.9668) | 0.9463 | 0.0663 | 0.0183 | 0.8577 | 0.9470 | 0.8805 | 0.8141 |
| Full AGVarPred | 120 | 0.42 | 0.9753 (0.9731-0.9775) | 0.9606 | 0.0568 | 0.0225 | 0.8938 | 0.9387 | 0.8945 | 0.8327 |

Note. Specificity recomputed as TN/(TN+FP) from confusion matrices; PR-AUC CIs were not computed; non-full models retrained with the regularized model's hyperparameters on the same splits.

**Supplementary Table S11.** AGVarPred performance stratified by gnomAD allele frequency

| **AF bin (raw gnomAD_AF)** | **N** | **Pathogenic** | **Benign** | **Prevalence** | **ROC-AUC (95% CI)** | **PR-AUC** | **F1** | **Sensitivity** | **Specificity** | **FPR** | **MCC** | **Brier** |
| --- | --- | --- | --- | --- | --- | --- | --- | --- | --- | --- | --- | --- |
| Absent (AF = 0) | 7,060 | 3650 | 3,410 | 0.5170 | 0.9373 (0.9318-0.9428) | 0.9460 | 0.8758 | 0.8701 | 0.8748 | 0.1252 | 0.7446 | 0.0934 |
| Ultra-rare (0 < AF < 10⁻⁴) | 959 | 873 | 86 | 0.9103 | 0.9821 (0.9727-0.9897) | 0.9980 | 0.9748 | 0.9966 | 0.5116 | 0.4884 | 0.6726 | 0.0334 |
| Rare (10⁻⁴ ≤ AF < 0.01) | 1,813 | 53 | 1,760 | 0.0292 | 0.9735 (0.9355-0.9992) | 0.9216 | 0.8224 | 0.8302 | 0.9943 | 0.0057 | 0.8171 | 0.0066 |
| Common (AF ≥ 0.01) | 2553 | 0 | 2,553 | 0.0000 | n/a | n/a | 0.0000 | 0.0000 | 1.0000 | 0.0000 | 0.0000 | 0.0001 |

Note. Bins mutually exclusive and exhaustive (sum = 12,385); AF = 0 denotes absence from gnomAD exomes r2.1.1; ultra-rare bin has only 86 benign variants and the rare bin only 53 pathogenic variants (small denominators).

**Supplementary Table S12.** Internal test-set performance by variant type

| **Variant type** | **N (%)** | **Pathogenic** | **Benign** | **Prevalence** | **ROC-AUC (95% CI)** | **PR-AUC** | **F1** | **Sensitivity** | **Specificity** | **MCC** | **Brier** |
| --- | --- | --- | --- | --- | --- | --- | --- | --- | --- | --- | --- |
| **SNV** | 9,773 (78.9%) | 2,240 | 7,533 | 0.2292 | 0.9558 (0.9519-0.9599) | 0.8892 | 0.7993 | 0.7866 | 0.9460 | 0.7410 | 0.0653 |
| **MNV** | 7 (0.06%) | 3 | 4 | 0.4286 | 1.0000 (1.0000-1.0000) | 1.0000 | 0.8000 | 0.6667 | 1.0000 | 0.7303 | 0.0795 |
| **Indel** | 2,605 (21.0%) | 2,333 | 272 | 0.8956 | 0.9532 (0.9336-0.9702) | 0.9909 | 0.9833 | 0.9970 | 0.7353 | 0.8280 | 0.0248 |

Note. SNV = single-nucleotide reference/alternate; MNV = equal-length substitution >1 nt; indel = unequal reference/alternate lengths. The MNV row (N = 7) is shown for completeness only and was excluded from interpretation. The indel subset is 89.6% pathogenic (ClinVar severity enrichment), which inflates PR-AUC/F1 relative to SNVs.

**Supplementary Table S13.** Internal and external calibration performance

| **Dataset** | **N** | **Prevalence** | **Brier** | **ECE (10 equal-width bins)** | **Calibration intercept** | **Calibration slope** |
| --- | --- | --- | --- | --- | --- | --- |
| Internal ClinVar test | 12,385 | 0.3695 | 0.0568 | 0.0225 | 0.115 | 0.898 |
| Humsavar | 14,718 | 0.0895 | 0.1828 | 0.1981 | -1.798 | 0.269 |
| Grimm2015 Selected | 5,253 | 0.0975 | 0.2691 | 0.2932 | -1.964 | 0.191 |
| DVD | 8,388 | 0.0232 | 0.1324 | 0.2182 | -5.613 | 1.509 |
| MAVE Independent | 7,557 | 0.4964 | 0.3935 | 0.3470 | 0.043 | 0.024 |
| VIP (consistency) | 3,880 | 0.2771 | 0.1920 | 0.2108 | -1.411 | 0.514 |
| gnomAD Common Benign | 4,989 | 0.0000 | n/a | n/a | n/a | n/a |
| PanelApp Benign | 1,773 | 0.0000 | n/a | n/a | n/a | n/a |
